# Data-efficient distal engineering of fluorinase using zero-shot models

**DOI:** 10.64898/2026.02.11.705267

**Authors:** David Harding-Larsen, Brianna M. Lax, Christopher Kurt Weingarten, Aboubakar Sako, Stanislav Mazurenko, Ditte Hededam Welner

**Affiliations:** The Novo Nordisk Foundation Center for Biosustainability, Technical University of Denmark, Søltofts Plads 220, 2200, Kongens Lyngby, Denmark; Loschmidt Laboratories, Department of Experimental Biology and RECETOX, Faculty of Science, Masaryk University, Kamenice 5, 625 00, Brno, Czech Republic; International Clinical Research Center, St. Anne’s University Hospital, Pekařská 664/53, 602 00, Brno, Czech Republic

## Abstract

Fluorinases have high potential for industrial biofluorination but any applications have been precluded by low catalytic efficiency and resistance to active site engineering. In this work, we employed PRIZM, a computational workflow utilizing an existing low-N dataset and zero-shot models for *in silico* prediction of activity-enhancing mutations at distal sites. The combination of these predictions with expert opinion led to the identification of 21 fluorinase mutants with enhanced relative activities, while 3 variants showed increased melting temperatures. A mutation in the hexameric interface, K237R, resulted in the largest stability gain, a more than 3.2-fold improvement in catalytic efficiency at 57°C, and an 8-fold increase in relative activity at 62°C. These results highlight the potential of distal fluorinase engineering for improving properties required to realize its industrial applications.

## INTRODUCTION

Fluorinases (5’-fluoro-5’-deoxyadenosine synthases, EC 2.5.1.63) are the only known class of enzymes able to directly catalyze carbon-fluorine bond formation, making them the subject of great interest for potential industrial bioproduction of organofluorides, which currently make up over 50% of agrochemicals^1,2^, 20% of marketed pharmaceuticals^3,4^, and contribute to many other sectors^5,6^. Fluorinases perform bimolecular nucleophilic substitution (S_N_2) on *S*-adenosyl-L-methionine (SAM) with fluoride as the nucleophile, producing 5’-fluoro-5’-deoxyadenosine (FDA) and L-methionine^7,8^ (Scheme 1). There are currently 18 functionally characterized fluorinases^9,10^, and two crystal structures available (fluorinases from *Streptomyces cattleya* and *Streptomyces* strain sp. MA37)^11,12^.

**Scheme 1.**
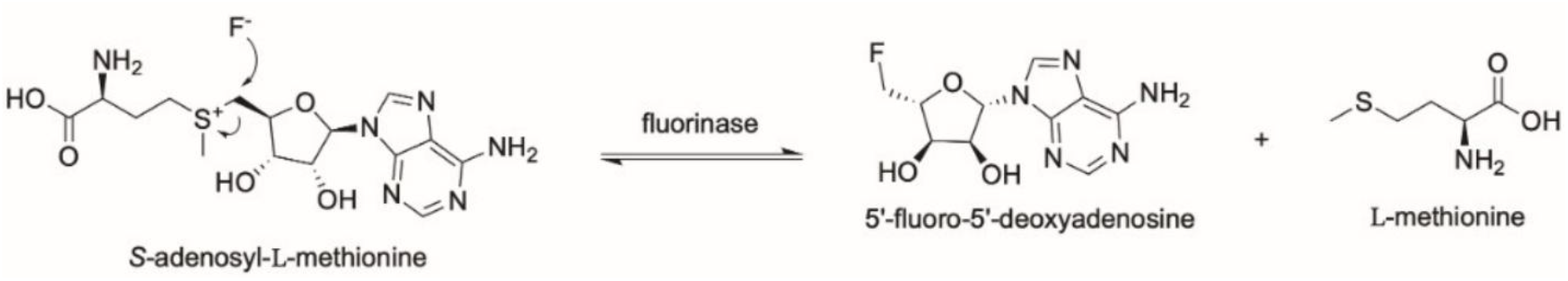
Reaction scheme of fluorinase-mediated SAM conversion into FDA and L-methionine.

Despite their potential, the fluorinases are infamous for their prohibitively slow turnover rates (*k*_*cat*_ = 0.16-0.41 min^−1^) and narrow substrate scopes that have precluded any industrial implementation^9^. Additionally, fluorinases have been historically resistant to catalytic improvements from active site mutagenesis, with previous active site engineering campaigns affording only incremental improvements^13,14^. One recent study performed rational engineering of the fluorinase at allosteric sites, and when combined with reaction medium optimization, was able to improve the turnover rate by 50x, representing a landmark improvement in biofluorination^15^. Thus, the existing literature suggests that the efficiency of fluorinases may perhaps be best enhanced outside of the active site.

Fluorinases belong to the “SAM-Dependent Hydrolases/Halogenases” superfamily, and despite similar monomeric structures to other members, they represent the only known hexameric halogenases^12,16^. The homohexameric structure is a dimer of trimers (D3 symmetry), containing a total of six identical monomers and six active sites. The monomer contains two domains, which are connected by a long loop present in all known fluorinases except for the archaeal fluorinase from *Methanosaeta* sp. PtaU1.Bin055^9^. The active sites are formed at each of the monomer interfaces in the trimers, containing residues from the N-terminal domain of one monomer and residues from the C-terminal domain of the other, obviating the need for trimerization in catalysis^12^. Our group recently showed that hexamerization, however, is not required for catalysis, although it does confer reaction efficiency by decreasing the Michaelis constant (*K*_*m*_) for SAM by two orders of magnitude^17^. The discovery of this functional benefit of hexamerization uncovered the hexameric interface as an interesting and unexplored target for engineering.

In recent years, protein engineering has seen the emergence of machine learning (ML) as a promising tool for guiding the mutagenesis process^18–20^. However, the performance of supervised models is tied to both the quantity and quality of datasets used for model training – properties that are often inversely related. An alternative approach is to employ these foundation models as zero-shot predictors, where the model outputs are used as proxies for fitness and activity changes caused by mutagenesis^21^. Since they do not rely on supervised learning, zero-shot predictors are versatile and simple to employ. Zero-shot predictors can be used across different scales – from engineering individual proteins^22–25^ to predicting variant effects at the proteome^26^, genome^27^, or even metagenome level^28^ – highlighting the versatility and broad biological knowledge contained in these pre-trained models.

In this study, we aimed to utilize ML to explore the distal mutational landscape in an intentional and efficient manner. First, we generated a low-N variant activity dataset of the fluorinase from *Streptomyces* sp. MA37 (hereafter referred to as “FlA”). We then employed our Protein Ranking using Informed Zero-shot Modelling (PRIZM) workflow, a two-phase pipeline, for identifying protein variants with enhanced properties^29^.

We expressed and characterized the best predicted single mutants, with ∼28% having higher activity than WT. We also explored combinations of mutations as well as semi-rational engineering informed by the ranked *in silico* library, both of which were successful. The best mutant K237R demonstrated a ∼3.6-fold improvement in catalytic efficiency at 57°C, as well as a ∼2.4°C increase in an apparent melting temperature (ΔT_m,app_).

Overall, this study shows that distal regions impact the activity of FlA and highlights the benefit of using ML to navigate these large sequence spaces. It also demonstrates the utility of zero-shot learning in engineering challenging, poorly understood enzymes such as FlA.

## RESULTS AND DISCUSSION

### Initial Experimental Characterization

We began our exploration of the FlA sequence space by generating a combinatorial mutant library of the hexameric interface residues. We targeted the hexameric interface for our mutational campaign since it is a unique feature of FlA within the halogenases and due to the limited success with active site engineering. Since the active sites are located at the monomer interfaces within the trimers, there is no overlap in residues that make up the active sites and those that participate in the hexameric interface. Based on our previous work, which demonstrated the hexameric region as a modulator of FlA activity^17^, we selected seven residues that facilitate this dimerization for our mutagenesis campaign (Figure 1A-B). The resulting library consisted of 1.28 billion variants, each containing mutations at between one and seven of these residues.

**Figure 1.**
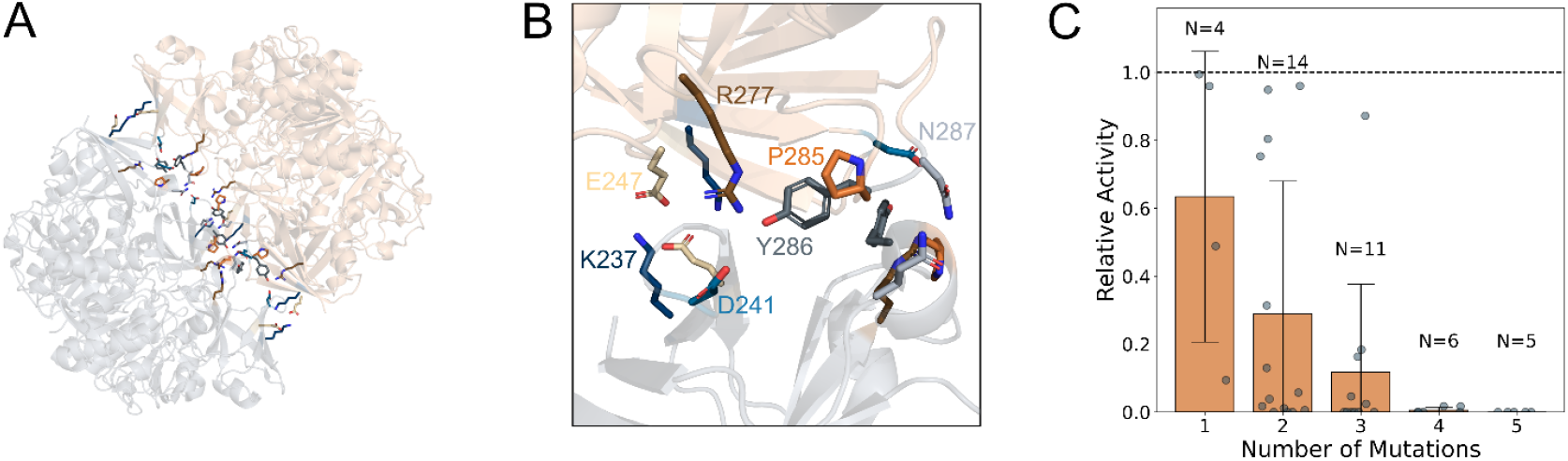
Initial FlA experimental examination. **A** Structural cartoon representation of the hexameric FlA (PDB ID: 5B6I), with each trimer distinctly colored and the six sets of seven hexameric interface residues shown as sticks. **B** Zoom-in on one of the contact points of the hexameric interface. **C** Relative activities for FlA variants in the low-N dataset as a function of the number of mutations per variant. Orange bars show average relative activity, grey circles indicate individual measurements, and error bars denote the standard deviation. The dashed line denotes WT activity of 1, with all variants being less active than WT.

Without a high-throughput screen available, it was unfeasible to characterize the entire library, so we randomly chose a small sample for initial characterization to look for trends between the mutation sites. Although we previously showed that hexamerization was not necessary for activity, the trimeric mutants that were tested contained only 1-2 mutations in the hexameric interface^17^. In this study, we characterized 40 variants with one to five mutations (Figure S1, Table S1). We observed an inverse relationship between the number of mutations per variant and relative activity, with the majority of variants with four or more mutations losing all activity (Figure 1C). Moreover, none of the tested variants had higher activity than WT, which prompted us to use ML to explore the mutational landscape *in silico* and narrow down promising variants for the expression and characterization in the lab.

### PRIZM Processing of FlA

Given the success of the PRIZM workflow at identifying beneficial non-active site mutants from small datasets in our previous work^29^, we decided to implement the workflow on our hexameric interface dataset. First, we used the 40 characterized variants to identify the best model for describing FlA relative activity (Figure 2A-C, Figure S2). ESM-IF1^30^, a structure-based zero-shot model, exhibited the best performance overall, while ESM-1b^27,31^ was the best performing sequence-based model, and TranceptEVE^22^ was the best performing model utilizing MSA information. We observed a significant difference in model performance between the best and worst models for our FlA dataset (Figure 2C), reinforcing that PRIZM can distinguish between the different zero-shot models using low-N datasets. While the ESM-IF1 scores were unable to accurately model the experimental data, a general trend was observed, where higher ESM-IF1 scores were associated with higher experimental values, in contrast to the low-scoring model UniRep. Notably, variants with no activity were very poorly predicted by both models, with inactive mutants giving rise to the full spectrum of scores. Comparing the ESM variants also highlighted that newer models do not always equal better performance, as the older ESM-1b outperformed the newer ESM-2^28^ (Figure S2).

**Figure 2.**
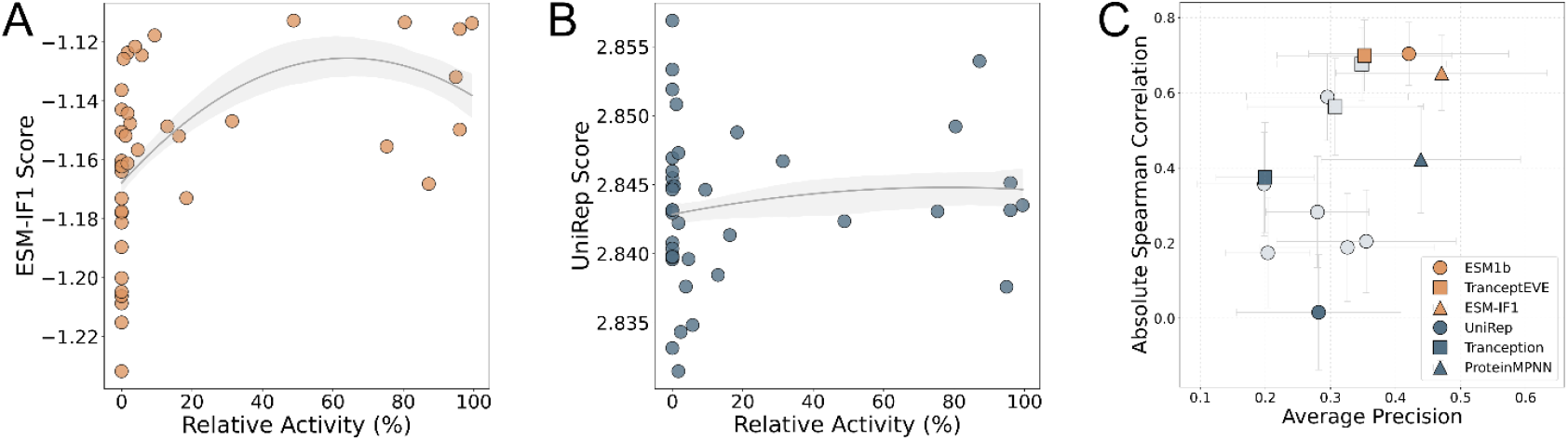
PRIZM model selection for FlA dataset. **A, B** Correlation between the experimental relative activities of the FlA low-N dataset and either ESM-IF1 scores (**A**) or UniRep scores (**B**). A second order polynomial is fitted to the data and is only used to guide the eye (the grey line denotes the regression estimate and the shaded area shows the confidence interval calculated using bootstrapping). **C** Spearman correlation and average precision for zero-shot scores and experimental values for the different zero-shot models of PRIZM (circle = sequence-based model, square = MSA-based model, and triangle = structure-based model; blue and orange denote the highest and lowest-correlated models for each model category, respectively, as identified by PRIZM). Error bars indicate the standard deviation estimated from bootstrapping the dataset (N = 1000).

We then constructed a full *in silico* FlA single-point mutant library and processed it using the top performers of each predictor category: ESM-1b (sequence-based), TranceptEVE (MSA-based), and ESM-IF1 (structure-based). This was partly due to the similar absolute Spearman correlations of the three models, but also because this allowed us to probe the effect of using sequence, evolutionary, and structural information for prediction. Using the ranked *in silico* library, we employed a *top N* approach for selecting the ten highest-ranked mutants for each zero-shot model. Furthermore, we found several mutants that were predicted to be better than WT by two of the models (Figure S3), suggesting that these variants are consistently favored by different zero-shot models trained on distinct types of protein information. As shown previously, combining model predictions from PRIZM can lead to enhanced variants^29^. We therefore also selected variants from model pairings – either by including all mutants predicted to be better than WT, or, when more than ten such mutants were identified, by combining the rankings from both zero-shot models and selecting the top candidates based on this combined ranking. Interestingly, no mutant scores were predicted to be better than WT by all three models at the same time. In total, 42 unique single mutants were chosen for experimental validation and named “the PRIZM variants” (Table S2).

### Experimental Validation of PRIZM Variants

We expressed the PRIZM variants recombinantly in *E. coli* and purified them using affinity chromatography. Two variants (P165N and F65I) did not express solubly and were omitted. For the remaining 40 variants, we examined the relative activity after two minutes (Figure S4) and three hours (Figure S5). Across all three top-performing zero-shot models, the activities of the selected variants were shifted toward higher values relative to the original low-N dataset, with many variants reaching or exceeding WT activity (Figure 3A). Interestingly, we observed significant variability in the distributions of the three selected zero-shot models; the variants chosen using ESM-1b scores were all worse than WT for both time points, while the variants of the structure-based ESM-IF1 had a large spread of activities, from almost no activity to higher than WT. The TranceptEVE variants performed the best on average, with the majority exhibiting enhanced activity compared to WT. These trends were mirrored in the variants selected by pairs of models, as the variants with ESM-1b being one of the selection models performed worse than those selected by TranceptEVE and ESM-IF1. The latter combination resulted in the best collection of variants, with a median relative activity of 1.085. These findings suggest that sequence alone might not be enough to guide engineering of FlA activity, and that epistatic effects captured in structural information could play a significant role in this enzyme. Interestingly, the relative activities converged around the WT value at the three-hour data point (Figure S6). We attributed this to product inhibition, a known limitation of fluorinases catalysis^15,32^, and continued with the two-minute data as the basis for identifying interesting mutations, due to the better separation of best and worst variants.

**Figure 3.**
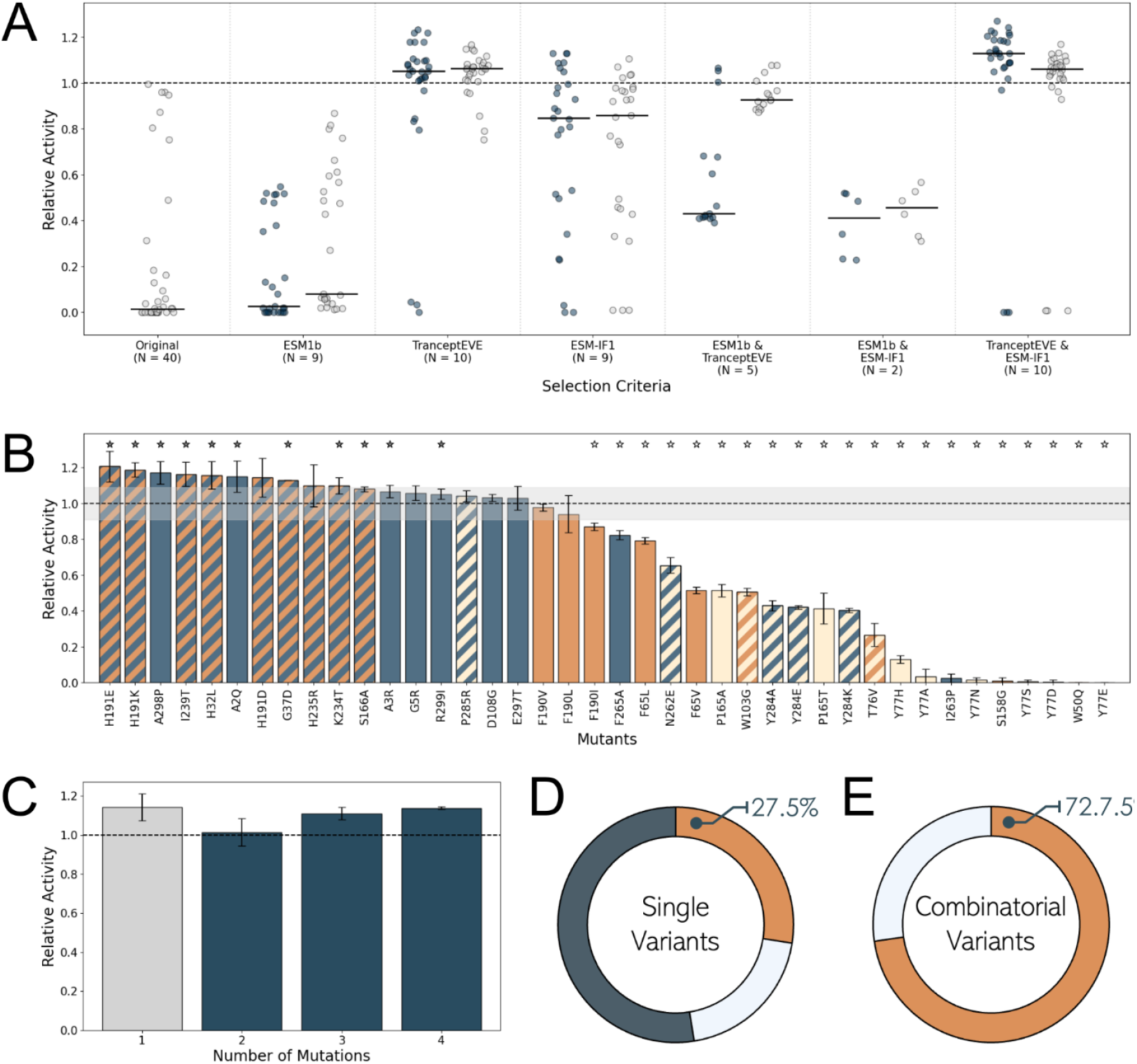
Relative activity of PRIZM mutants. Dashed lines denote the WT activity. **A** Point clouds of relative activities after two minutes (blue) and three hours (grey) for variants selected using different criteria. “Original” denotes the original low-N FlA dataset. Horizontal lines denote median. **B** Relative activities after two minutes of single mutants from different selection criteria: beige = ESM-1b, blue = TranceptEVE, and orange = ESM-IF1. Striped colors denote variants chosen based on two models. Stars denote variants with relative activity at two minutes significantly different from WT (one-sided Welch test with α = 0.05, grey = better than WT, white = worse than WT). Although certain variants are selected based on multiple selection criteria, each is displayed only once in this figure (for a comprehensive overview of all selections, refer to Figure S4). Error bars indicate the standard deviation of biological replicates. **C** Average relative activities after two minutes as a function of the number of mutations per variant. Grey = selected single variants, blue = combinatorial variants. Error bars indicate the standard deviation, while the relative activities of individual combinatorial mutants can be found in Figure S7. Error bars indicate the standard deviation of biological replicates. **D, E** Percentage of variants with significantly higher activity than WT (orange), WT-like activity (light blue), or lower activity than WT (dark grey) at two minutes for PRIZM/single (**D**) and combinatorial (**E**) variants. Significance was determined using a one-sided Welch test (α = 0.05).

We observed 11 variants with activities significantly above WT (Figure 3B), with the best variant, H191E, having a relative activity of 120.7%. To explore potential epistasis, from these 11 variants, we selected four mutations for combinatorial analysis: H191E, H32L, G37D, and S166A. These variants were selected since they were the highest-ranked variants with unique positions from the top-performing PRIZM result (combination of TranceptEVE and ESM-IF1, Table S2). Interestingly, S166A was previously reported to be important for enzyme activity for the fluorinase from *Amycolatopsis* sp. CA-128772 due to its influence on the fluoride ion channel^14^. This study analyzed 62 mutants to identify this mutation, whereas it was ranked as the tenth best by ESM-IF1 and third best by the combination of TranceptEVE and ESM-IF1, significantly reducing the number of variants tested to arrive at this interesting mutation. Lastly, it is worth noting that all the active site mutations predicted to be better than WT exhibited little activity, confirming the challenge of engineering the active site of fluorinases.

We combined these four single mutants into double, triple, and quadruple mutants, denoted as “the combinatorial variants” (Figure 3C, Figure S7). Here, most of the double mutants exhibited no enhanced performance compared to WT, with only H32L-H191E having similar performance to the corresponding single mutant variants. The triple and quadruple mutants recovered the activity levels of the single mutant variants but did not exhibit additional synergy. Interestingly, we did not observe the same trend as the original low-N dataset, where an increase in the number of mutations was inversely correlated with the relative activity of FlA (Figure 1C).

In conclusion, combining PRIZM with a greedy approach resulted in a best-performing single mutant (H191E) with 120.7% relative activity. H191 is located within the linker connecting the two domains of the FlA monomer (Figure S7), a site that is spatially distant from conventionally studied regions. As such, H191E would likely have been overlooked in traditional rational engineering approaches. This finding illustrates the utility of zero-shot learning methods in identifying functionally relevant mutations in regions of the protein landscape that lack prior functional annotation or intuitive significance. Overall, we had a success rate for variants with higher activity than WT of 27.5% (11/40) for the PRIZM/single mutants and 50% (3/6) for the double mutants, while all the triple and quadruple mutants had activity above the WT, resulting in an overall 72.7% success rate for the combinatorial variants (Figure 3D-E).

### Targeted Engineering using PRIZM and MSA

Following our PRIZM and combinatorial variants, we tested a more targeted engineering approach, hypothesizing that combining expert knowledge with zero-shot prediction would enhance success rate and variant performance; Our experimentally validated PRIZM variants with enhanced activity located primarily to three regions of interest (Figure 4A). The first is the hexameric interface, *i*.*e*., the original target of our low-N dataset. The two others are the N-terminal tunnel and the area around the active site (henceforth denoted as the second-shell of the active site). We utilized the PRIZM rankings to identify residues within these regions predicted to be better than WT by both TranceptEVE and ESM-IF1. To complement the PRIZM analysis, we constructed an MSA of previously studied fluorinase homologues^9^ (Table S3). This homologue MSA was utilized to select mutations at our identified positions that were either conserved across multiple fluorinases (consensus) or present in the faster fluorinase homologues.

**Figure 4.**
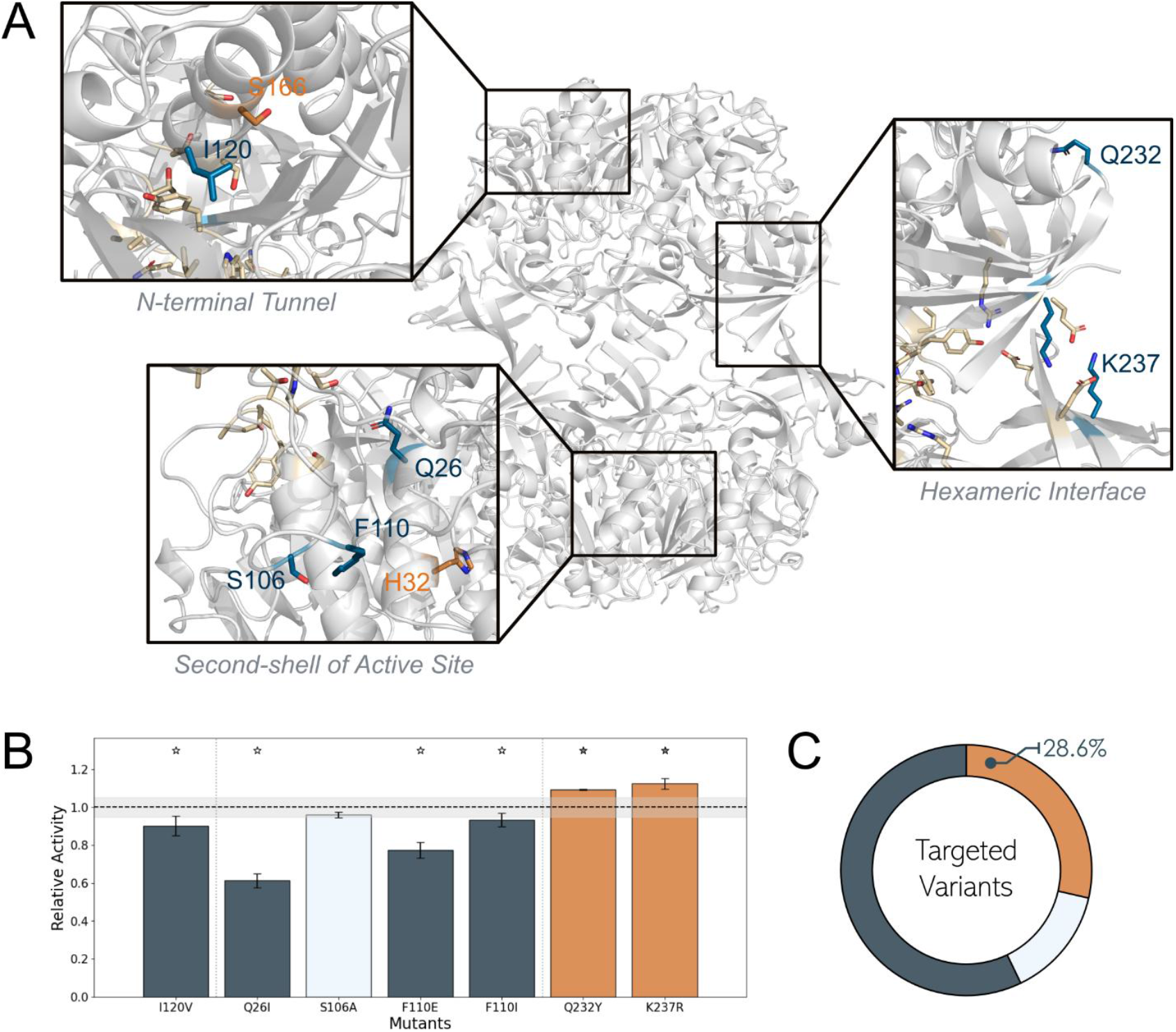
Targeted engineering of FlA. **A** Close-up of regions of interest used for targeted engineering, with residues of note shown as sticks (PDB ID: 5B6I). Newly selected residues are colored blue, previous PRIZM variants are colored orange, while active site residues or initial hexameric interface residues are colored beige. **B** Relative activity of targeted variants at two minutes. Variants with significantly higher activity than WT are colored orange, WT-like variants are colored light blue, while those with lower activity than WT are dark grey. Dashed line denotes the WT value of 1. Error bars indicate the standard deviation of biological replicates with N = 3. Stars denote variants with relative activity at two minutes significantly different from WT (one-sided Welch test with α = 0.05, grey = better than WT, white = worse than WT) are denoted by a star. **C** Percentage of variants with significantly higher activity than WT (orange), WT-like activity (light blue), or lower activity than WT (dark grey) at two minutes for targeted variants. Significance was determined using a one-sided Welch test (α = 0.05).

In the hexameric interface, we identified Q232Y and K237R, with both mutations being highly conserved across the other fluorinase homologues. For the N-terminal tunnel, we selected I120V, as the MSA showed the valine to be present at this position in several of the homologues, including two of the faster fluorinases. Lastly, four mutations were chosen at the second shell of the active site: Q26I, with the isoleucine being present in the fastest homologue, F110I and F110E, where the mutated residue was present across multiple homologues, as well as S106A. In total, we selected and characterized seven additional mutations, denoted as the “targeted variants”.

Many of these targeted variants showed either close to or worse activity than that of the WT enzyme (Figure 4B). Especially the Q26I variant, situated close to the active site, showed decreased performance, consistent with previous challenges of improving the enzyme with mutations near the active site. However, both mutations in the hexameric interface exhibit significantly enhanced performance, confirming our initial hypothesis of the importance of this region for FlA activity. Overall, our targeted approach achieved a success rate of 28.6% (2/7), still outperforming our initial greedy approach and highlighting strength of combining expert knowledge with the versatility and power of the raw zero-shot predictions (Figure 4C).

### Biochemical Characterization of Top Variants

We then investigated the underlying causes of the increased relative activities in our top variants: the four previously selected PRIZM variants, H32L, G37D, S166A, and H191E, as well as the best targeted mutant, K237R.

We first quantified the T_m,app_ of each of these variants using differential scanning fluorimetry (DSF), in reaction buffer alone, with SAM, or with F^−^ (Figure 5A, Figure S9). We observed that the T_m,app_ values of H191E, H32L, and K237R were all increased compared to WT in all three conditions, with the latter having a ΔT_m,app_ value of up to ∼2.4°C in the presence of SAM. In particular, K237R exhibited an increase in ΔT_m,app_ in the presence of the substrates, suggesting a synergy between the mutation and the interaction of the protein with these molecules. We see that all mutants except S166A and all except G37D and S166A have larger increases in ΔT_m,app_ after the binding of F^−^ and SAM, respectively, compared to WT (Figure S9).

**Figure 5.**
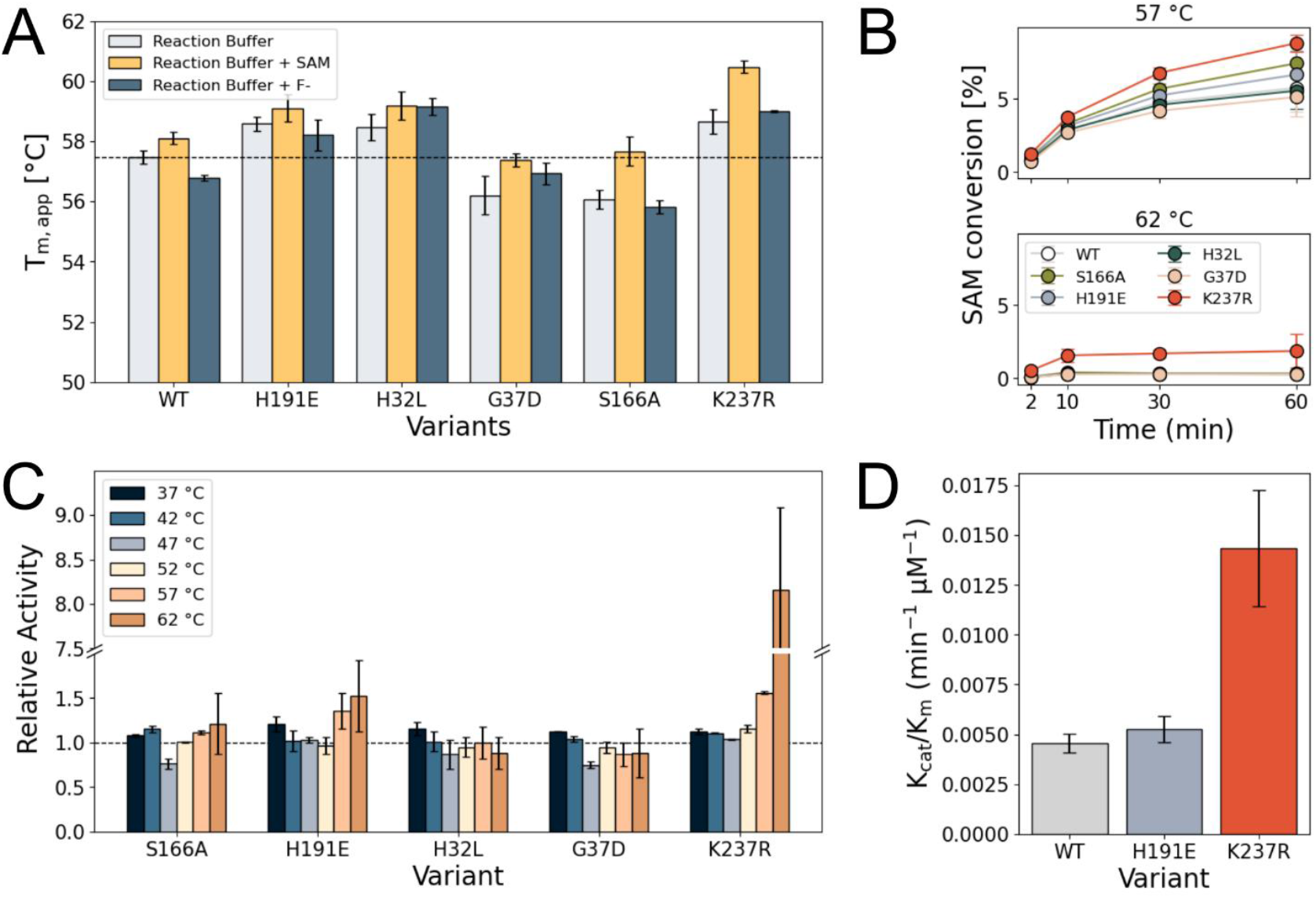
Biochemical characterization of FlA WT and top FlA variants. Error bars indicate the standard deviation of biological replicates with N = 3. **A** T_m,app_ in reaction buffer alone (grey), with SAM (yellow), or with F^−^ (blue). The dashed line indicates the T_m,app_ of WT in the reaction buffer. **B** SAM conversion at 57°C (top) and 62°C (bottom) at 2, 10, 30, and 60 minutes. **C** Relative activity after two minutes at different temperatures, with the dashed lined indicating a relative activity of 1. **D** Catalytic efficiency at 57°C.

To complement our thermostability assay, we examined the activity of the variants at 2, 10, 30, and 60 minutes at temperatures ranging from 37°C to 62°C (Figure 5B, Figure S10-11). We observed an optimal temperature of WT around 52°C, with the enzyme appearing mostly inactive at 62°C, which aligns with this being above the enzyme’s T_m,app_. Despite the increases in T_m,app_ values for many of the mutants, the optimal temperature remained around 52°C for all variants. Interestingly, while S166A, H191E, and K237R demonstrated enhanced relative activity at 37°C, they outperformed WT to a much greater extent at 57°C and 62°C (Figure 5C, Figure S12). The three variants achieved a 21.4%, 52.6%, and 716.2% increase in relative activity, respectively, at 62°C. Given these impressive increases in relative activities for H191E and K237R at elevated temperatures, we performed Michaelis-Menten kinetic analysis at 57°C (Figure 5D, Table 1, Figure S14). We did not observe an increase in the turnover rate of either of these variants, but rather, an improvement in catalytic efficiency related to decreased Michaelis constants (*K*_*m*_) for SAM. The *K*_*m*_ values for H191E and K237R were ∼1.2-fold and ∼3.6-fold lower than WT, respectively, leading to 18% and ∼218% increases in catalytic efficiency. For K237R, this is in alignment with our data showing that the greatest stability benefits from these mutations are observed in the presence of SAM. We believe that these mutations are able to improve SAM binding, which then improves the stability of the protein, synergistically improving the efficiency of the enzyme without affecting the turnover rate.

**Table 1.** Biochemical characterization of WT, H191E, and K237R. Error is based on the standard deviation of biological replicates with N = 3.

| Variant | $T_{m,app}$ (alone)<br>°C | $T_{m,app}$ (SAM)<br>°C | $T_{m,app}$ (F<sup>-</sup>)<br>°C | $k_{cat}$<br>min<sup>-1</sup> | $K_m$<br>μM | Catalytic efficiency<br>min<sup>-1</sup> μM<sup>-1</sup> |
| --- | --- | --- | --- | --- | --- | --- |
| WT | 57.5 ± 0.2 | 58.1 ± 0.2 | 56.8 ± 0.1 | 1.10 ± 0.13 | 244 ± 39 | 0.0045 ± 0.0005 |
| H191E | 58.6 ± 0.2 | 59.1 ± 0.5 | 58.2 ± 0.5 | 1.09 ± 0.12 | 209 ± 43 | 0.0053 ± 0.0007 |
| K237R | 58.7 ± 0.4 | 60.5 ± 0.2 | 59.00 ± 0.02 | 0.94 ± 0.07 | 67 ± 15 | 0.0143 ± 0.0029 |

Based on this biochemical characterization, we hypothesize that the improved performances of the H191E and K237R variants arise primarily from an increase in SAM affinity, which provides stabilization to the enzyme and the active sites. These mutations do not inherently increase the catalytic rate of the enzyme but rather enable a more efficient binding of SAM within the active site. This is particularly important for fluorinases because it is known that SAM binding is required for the full desolvation of the bound fluoride ions, which is the most energetically expensive part of the reaction^32^. An increased affinity for SAM can help to address this constraint.

H191 is located in the linker region between the two domains of the FlA monomer (Figure S8). Substitutions with glutamate could perturb the microenvironment, although the increased relative activity observed for H191K (Figure 3B) suggests that the effect is not dependent on the charge sign. Instead, introducing a charge at this position may facilitate better solvation of the linker, reduce hydrophobic frustration in this region, and stabilize the fold. This could ultimately lead to more stable active sites, which can better bind to SAM.

K237 is instead situated in the hexameric interface across from E247 (Figure S13). Here, the arginine substitution likely enhances the interactions with E247, either by strengthening existing hydrogen bonds or forming additional electrostatic contacts, thereby stabilizing the hexameric state of FlA. This is in agreement with our previous work that showed a trimeric FlA suffering a 200-fold increase in SAM *K*_*m*_ compared to the hexameric WT^17^. Together with this work, these results highlight the catalytic relevance of this unique structural feature of fluorinases.

Finally, we were unable to elucidate the mechanisms of increased relative activity for H32L, G37D, and S166A. The last mutation was previously suggested to improve fluoride entry through the N-terminal tunnel^14^, which we did not investigate here. For H32L and G37D, the effect may instead relate to improved resilience toward product inhibition, an influence highlighted by Slanska et al.^15^, though such a characterization is outside of the scope of this study.

## CONCLUSION

In this study, we identified several FlA variants with enhanced performance using PRIZM, a novel zero-shot prediction-based approach for protein engineering. Using only PRIZM predictions, we achieved a 27.5% hit rate of single mutants with relative activity above WT. A combinatorial screen of the best variants resulted in a hit rate of 72.7%, while a targeted approach exhibited a 28.5% hit rate. Importantly, PRIZM is simple and versatile as it only needs a low-N dataset and does not require any supervised learning. Moreover, despite the original low-N dataset only having mutations in the hexameric interface, we were able to identify variants with increased activity across the entire protein sequence.

Our best PRIZM variant, H191E, and our best targeted variant, K237R, demonstrated 52.6% and 716.2% increases in relative activity at 62°C, respectively. Our biochemical characterization indicated that this is a result of stabilization of the enzyme and enhanced SAM binding; Given that the highest increase in T_m,app_, ∼2.4°C, was observed in the presence of SAM, we hypothesized that the mutations are not only stabilizing, but also improve binding to SAM, which then also contributes to the overall enzyme stability. This hypothesis was confirmed via kinetic analysis at elevated temperatures, which revealed a ∼3.6-fold decrease in SAM *K*_*m*_ and a ∼218% increase in catalytic efficiency for K27R. For H191E, we hypothesize that introducing a charge in the linker region could improve solvation and reduce hydrophobic frustration.

Our analysis confirms the potential of the hexameric interface for modulating FlA activity. In addition, this study identified new distal regions important for FlA performance that do not directly interfere with the active site, such as the linker region between the two domains of the monomer and the second shell of the active site. Future exploration of these regions may provide insight into the mechanisms underlying the distal effects on FlA activity, with potential value for engineering FlA to enhance industrial performance.

## MATERIALS AND METHODS

### Fluorinase Data Setup

The FlA low-N data was pre-processed by omitting all variants showing inconclusive relative activity. The structural model was created using AlphaFold3^33^ while the MSA was obtained using the EVcouplings alignment module ^34^ with a bitscore cutoff of 0.3. The binarization threshold was chosen to be a relative activity of 80% due to no variant exhibiting higher-than-wildtype activity. The *in silico* library of FlA contained all single-point mutants for the entire protein sequence and was created using our PRIZM pipeline^29^.

### Materials

5’-fluoro-5’-deoxyadenosine (FDA) and 5’-chloro-5’-deoxyadenosine (ClDA) were kindly provided to us by Eray Bozkurt (Pablo Nikel, DTU Biosustain) and stored at −20°C. All other materials used in this work are commercially available.

### Library Construction

The DNA library of hexameric interface mutants of the FlA gene from *Streptomyces* sp. strain MA37 (FlA^MA37^) was ordered from Neochromosome. The library was generated using saturation mutagenesis at the following seven hexameric interface residues: K237, D241, E247, R277, P285, Y286, and N287. The library was amplified via PCR and cloned into the pET28a(+) backbone for protein expression using USER cloning^35^. The USER reaction mixtures were transformed into *E. coli* and the mutants were purified as described in the following sections.

### Cloning & Transformation

The WT FlA gene from *Streptomyces* sp. strain MA37 (FlA^MA37^) was previously cloned into the pET28a(+) backbone with an N-terminal His-tag and tobacco etch virus (TEV) cleavage site^17^. Primers were designed using NEBaseChanger^36^ and introduced into the FlA gene with site directed mutagenesis via PCR, followed by incubation with the Kinase, Ligase, and DpnI (KLD) Enzyme Mix (New England Biolabs) for 1h at room temperature (RT). The KLD reaction mixtures were then transformed into chemically competent DH5α *E. coli* cells (New England Biolabs) via heat shocking for 30s at 42°C, recovered in Super Optimal broth with Catabolite repression (SOC) medium for 1h and then plated on agar plates containing kanamycin (50 µg/mL) overnight. Individual colonies were picked and grown in liquid cultures, miniprepped (Macherey-Nagel), and the resulting plasmids were sequenced via Sanger sequencing with T7 promoter and terminator primers to confirm the introduction of the desired mutation (Eurofins).

### Protein Expression & Purification

Plasmids with confirmed sequences were transformed via heat shock into BL21(DE3) (Invitrogen) chemically competent *E. coli* as stated above. Expression cultures were inoculated 1:1000 with the overnight pre-cultures and grown at 37°C to an optimal density at 600 nm (OD_600_) of ∼0.8. Cultures were induced with 250 µM isopropyl β-?-1-thiogalactopyranoside (IPTG) and incubated overnight (at least 20h) at 18°C. Cells were harvested by centrifugation and either directly used in purification or flash frozen in liquid nitrogen and stored at −20°C until purification. All further steps during purification were performed either on ice or at 4°C in a cold room. Pellets were resuspended in 15 mL Lysis Buffer (20 mM imidazole, 300 mM NaCl, 10 mM 4-(2-Hydroxyethyl)piperazine-1-ethane-sulfonic acid (HEPES), 0.05% (v/v) Triton X-100, 10 µg/mL DNase, pH 7.4) and lysed via sonication for 3min (15s on/30s off) with 60% amplitude with a 6-mm probe (Sonics & Materials Inc.). The lysate was then centrifuged and filtered.

The samples were purified using an ÄKTA Pure M fast protein liquid chromatography (FPLC) system (Cytiva Life Sciences) with a 1 mL HisTrap FF column (Cytiva Life Sciences). The samples were loaded on the column, washed with 20 mL 90% Buffer A (20 mM imidazole, 300 mM NaCl, 10 mM HEPES, pH 7.4) and 10% Buffer B (500 mM imidazole, 300 mM NaCl, 10 mM HEPES, pH 7.4), and eluted with 15 mL 40% Buffer A and 60% Buffer B into 1 mL fractions. Protein molecular weight (MW) and purity was characterized using SDS-PAGE on a NuPAGE 4-12% Bis-Tris polyacrylamide gel (Invitrogen). Peak fractions were pooled and buffer exchanged into Buffer C (150 mM NaCl, 50 mM HEPES, pH 7.4) using PD-10 desalting columns (Cytiva Life Sciences). Purified proteins were concentrated to a final concentration of 5 mg/mL using 30 kDa Amicon Ultra Centrifugal Filters (Merck Millipore), flash frozen in liquid nitrogen, and stored at −70°C for further use.

### Enzyme Activity Assays

Protein aliquots were thawed on ice and the concentrations were measured via NanoDrop (Thermo Scientific) before diluting all samples to 2 mg/mL. A separate master mix was made for each replicate, with a final substrate concentration of 300 µM SAM and 75 mM KF in Reaction Buffer (150 mM NaCl, 10 mM HEPES, pH 7.8). The master mixes were aliquoted in PCR tube strips, and the protein variants were added simultaneously with a multichannel pipette to a final hexamer concentration of 1 µM, based on the hexameric molecular weight (204 kDa). The reactions were incubated at 37°C (unless otherwise stated) and quenched via heat inactivation at 95°C for 5min. For the relative activity assays, the 3h timepoint, the protein was added to the master mix on ice before being transferred to a thermocycler, while the protein was added in the pre-heated samples for the 2min timepoint. The samples were diluted 2× and filtered using 0.45-µm 96-well plate Durapore membranes (Millipore) and run on the HPLC. For quantification of SAM conversion, the FDA concentration was divided by the total amount of FDA and SAM (FDA concentration and SAM degradation product concentration).

For determining the Michaelis-Menten parameters of the variants, reactions were performed using 1 µM of enzyme, based on the hexameric molecular weight, 10-1200 µM SAM, and 75 mM KF in Reaction Buffer. The reactions were performed at 57°C for 15s and 30s and quenched via heat inactivation at 95°C for 5min. Three replicates were performed on separate days. Michaelis-Menten kinetics were assumed to determine *k*_*cat*_ and *K*_*M*_. The velocity of the enzyme was determined for each SAM concentration using linear regression and the Michaelis-Menten equation was then fit to the velocity-concentration dataset using GraphPad Prism (version 10.6.1).

### High Performance Liquid Chromatography (HPLC)

The reaction compositions were analyzed by Reversed-Phase HPLC using an Ultimate 3000 Series HPLC (Thermo Fisher Scientific) and a ZORBAX RR Eclipse Plus C18 Column (100 × 4.6 mm, 3.5 µm) (Agilent) maintained at 30°C. MilliQ water + 0.1% (v/v) formic acid and acetonitrile were used as mobile phases A and B, respectively, with the following method in percentages of mobile phase B at 1 mL/min: 0-1.5 min 5-12% (gradient), 1.5-2.5 min 12%, 2.5-4.5 min 12-30% (gradient), 4.5-5 min 30-70% (gradient), 5-5.5 min 70-5% (gradient), 5.5-7 min 5%. Chromatograms were recorded at 260 nm and integrated using the Chromeleon software (version 7.2.9). Using this method, FDA elutes at 2.64 min, a SAM degradation product elutes at 3.81 min, and vanillin elutes at 6.35 min (Figure S15). Standard curves correlating FDA and SAM concentrations with the ratios of their absorbances at 260 nm were generated using solutions of FDA or SAM from 0.5-30 µM and .5-300 µM, respectively (Figure S16). SAM cannot be measured directly on the HPLC due to its instability, so the standard curve was created using the degradation product mentioned above.

### Differential Scanning Fluorimetry (DSF)

The melting temperatures of the purified enzymes were determined using the Protein Thermal Shift Dye Kit (Thermo Fisher Scientific) and a qPCR QuantStudio 5 Real-Time PCR machine (Applied Biosystems). Immediately before the experiment, 50 µL of 2× dye was mixed with 50 µL of enzyme at 1 mg/mL. The melting temperatures were measured in the following conditions: (1) in Reaction Buffer, (2) in Reaction Buffer with 75 mM KF, and (3) in Reaction Buffer with 300 µM SAM. The samples were incubated at 25°C for 2min, then the temperature was linearly increased to 99°C. All data analysis was performed using Protein Thermal Shift Software v1.4.

## Supporting information

FlA variant zero-shot scores and experimental results

## ASSOCIATED CONTENT

### Supporting Information

Supporting Information: Correlation between zero-shot scores of FlA *in silico* library from best models; FlA mutational landscape; structural illustrations of FlA; relative activity measurements of single and combinatorial variants; Comparison of WT and top FlA variants (ΔT_m,app_, temperature optimum, relative activity at different temperatures after 2 minutes and 10 minutes; representative Michaelis Menten curve, representative HPLC trace; SAM and FDA calibration curves; and supporting datasets including variant libraries and multiple sequence alignments (PDF). Zero-shot scores of *Gm*SuSy single mutants and experimental results for top *Gm*SuSy variants (XLSX). Zero-shot scores of TOGT1_1 single mutants and experimental results for selected TOGT1_1 variants (XLSX).

### Data Availability Statement

All data, analysis scripts, and code related to PRIZM used in this study are openly available at GitHub: https://github.com/daha-la/PRIZM. The version of the PRIZM workflow used in this study corresponds to the GitHub release v1.0.0, which contains the complete codebase, analysis scripts for reproducing the figures, and the datasets underlying the study.

## AUTHOR INFORMATION

### Author Contributions

The manuscript was written through the contributions of all authors. All authors have given approval to the final version of the manuscript.

### Funding Sources

We gratefully acknowledge financial support from the Novo Nordisk Foundation (grant NNF20CC0035580). Additional funding was provided by the Czech Ministry of Education, Youth and Sports (ESFRI RECETOX RI LM2023069, ESFRI ELIXIR LM2023055, and e-INFRA CZ 90254) as well as the European Union’s Horizon 2020 Research and Innovation Programme through grant agreement No. 857560 (CETOCOEN Excellence) and the European Union’s Horizon Europe Framework Programme under the grant agreement No. 101136607 (CLARA). This publication reflects only the author’s view, and the European Commission is not responsible for any use that may be made of the information it contains.

### Notes

The authors report no conflicts of interest.

## ACKNOWLEDGMENT

The authors thank Eray Bozkurt (Pablo Nikel, DTU Biosustain) for kindly providing us with FDA and ClDA.

## ABBREVIATIONS

ClDA: 5’-chloro-5’-deoxyadenosine
FDA: 5’-fluoro-5’-deoxyadenosine
FlA: fluorinase from *Streptomyces* sp. MA37
HPLC: high-performance liquid chromatography
ML: machine learning
PDB ID: Protein Data Bank Identifier
PRIZM: Protein Ranking using Informed Zero-shot Modelling
SAM: S-adenosyl-L-methionine
WT: wildtype.

## Notes

### Competing Interest Statement

The authors have declared no competing interest.

## REFERENCES

(1) Ogawa, Y.; Tokunaga, E.; Kobayashi, O.; Hirai, K.; Shibata, N. Current Contributions of Organofluorine Compounds to the Agrochemical Industry. iScience 2020, 23 (9), 101467. 10.1016/J.ISCI.2020.101467.

(2) Martinelli, L.; Nikel, P. I. Breaking the State-of-the-Art in the Chemical Industry with New-to-Nature Products via Synthetic Microbiology. Microb. Biotechnol. 2019, 12 (2), 187–190. 10.1111/1751-7915.13372.

(3) Inoue, M.; Sumii, Y.; Shibata, N. Contribution of Organofluorine Compounds to Pharmaceuticals. ACS Omega 2020, 5 (19), 10633–10640. 10.1021/ACSOMEGA.0C00830.

(4) Chandra, G.; Singh, D. V.; Mahato, G. K.; Patel, S. Fluorine-a Small Magic Bullet Atom in the Drug Development: Perspective to FDA Approved and COVID-19 Recommended Drugs. Chemical Papers 2023 77:8 2023, 77 (8), 4085–4106. 10.1007/S11696-023-02804-5.

(5) Zhang, W.; Liang, Y. The Wide Presence of Fluorinated Compounds in Common Chemical Products and the Environment: A Review. Environmental Science and Pollution Research 2023, 30 (50), 108393–108410. 10.1007/S11356-023-30033-6,.

(6) Zhang, C.; Yan, K.; Fu, C.; Peng, H.; Hawker, C. J.; Whittaker, A. K. Biological Utility of Fluorinated Compounds: From Materials Design to Molecular Imaging, Therapeutics and Environmental Remediation. Chem. Rev. 2022, 122 (1), 167–208. 10.1021/ACS.CHEMREV.1C00632.

(7) Cadicamo, C. D.; Courtieu, J.; Deng, H.; Meddour, A.; O’Hagan, D. Enzymatic Fluorination in Streptomyces Cattleya Takes Place with an Inversion of Configuration Consistent with an SN2 Reaction Mechanism. ChemBioChem 2004, 5 (5), 685–690. 10.1002/CBIC.200300839.

(8) O’Hagan, D.; Schaffrath, C.; Cobb, S. L.; Hamilton, J. T. G.; Murphy, C. D. Biosynthesis of an Organofluorine Molecule: A Fluorinase Enzyme Has Been Discovered That Catalyses Carbon-Fluorine Bond Formation. Nature 2002, 416 (6878), 279. 10.1038/416279A.

(9) Pardo, I.; Bednar, D.; Calero, P.; Volke, D. C.; Damborský, J.; Nikel, P. I. A Nonconventional Archaeal Fluorinase Identified by in Silico Mining for Enhanced Fluorine Biocatalysis. ACS Catal. 2022, 12 (11), 6570–6577. 10.1021/ACSCATAL.2C01184.

(10) Verma, R. K.; Yeo, W. L.; Tiong, E.; Ang, E. L.; Lim, Y. H.; Wong, F. T.; Fan, H. Unveiling the Molecular Basis of Selective Fluorination of SAM-Dependent Fluorinases. Chem. Sci. 2025, 16 (23), 10610–10619. 10.1039/D5SC00081E.

(11) Deng, H.; Ma, L.; Bandaranayaka, N.; Qin, Z.; Mann, G.; Kyeremeh, K.; Yu, Y.; Shepherd, T.; Naismith, J. H.; O’Hagan, D. Identification of Fluorinases from Streptomyces Sp MA37, Norcardia Brasiliensis, and Actinoplanes Sp N902-109 by Genome Mining. ChemBioChem 2014, 15 (3), 364–368. 10.1002/CBIC.201300732.

(12) Dong, C.; Huang, F.; Deng, H.; Schaffrath, C.; Spencer, J. B.; O’Hagan, D.; Naismith, J. H. Crystal Structure and Mechanism of a Bacterial Fluorinating Enzyme. Nature 2004, 427 (6974), 561–565. 10.1038/NATURE02280.

(13) Sun, H.; Yeo, W. L.; Lim, Y. H.; Chew, X.; Smith, D. J.; Xue, B.; Chan, K. P.; Robinson, R. C.; Robins, E. G.; Zhao, H.; Ang, E. L. Directed Evolution of a Fluorinase for Improved Fluorination Efficiency with a Non-Native Substrate. Angewandte Chemie - International Edition 2016, 55 (46), 14277–14280. 10.1002/ANIE.201606722.

(14) Feng, X.; Cao, Y.; Liu, W.; Xian, M. Identification of Two Novel Fluorinases From Amycolatopsis Sp. CA-128772 and Methanosaeta Sp. PtaU1.Bin055 and a Mutant With Improved Catalytic Efficiency With Native Substrate. Front. Bioeng. Biotechnol. 2022, 10, 881326. 10.3389/FBIOE.2022.881326.

(15) Slanska, M.; Volke, D. C.; Pardo, I.; Kunka, A.; Krishna, N. B.; Shetty, A. J.; Muthuraj, L.; Sigamani, G.; Lalitha, R.; Büll, A. K.; Marek, M.; Kumar, P. R.; Damborsky, J.; Nikel, P. I.; Prokop, Z. Mechanism-Guided Engineering of Fluorinase Unlocks Efficient Nucleophilic Biofluorination. bioRxiv 2025, 2025.07.28.666932. 10.1101/2025.07.28.666932.

(16) Eustáquio, A. S.; Pojer, F.; Noel, J. P.; Moore, B. S. Discovery and Characterization of a Marine Bacterial SAM-Dependent Chlorinase. Nat. Chem. Biol. 2008, 4 (1), 69–74. 10.1038/NCHEMBIO.2007.56.

(17) Kittilä, T.; Calero, P.; Fredslund, F.; Lowe, P. T.; Tezé, D.; Nieto-Domínguez, M.; O’Hagan, D.; Nikel, P. I.; Welner, D. H. Oligomerization Engineering of the Fluorinase Enzyme Leads to an Active Trimer That Supports Synthesis of Fluorometabolites in Vitro. Microb. Biotechnol. 2022, 15 (5), 1622–1632. 10.1111/1751-7915.14009.

(18) Markus, B.; Christian C G.;, Andreas, K.; Arkadij, K.; Stefan, L.; Gustav, O.; Elina, S.; Radka, S. Accelerating Biocatalysis Discovery with Machine Learning: A Paradigm Shift in Enzyme Engineering, Discovery, and Design. ACS Catal. 2023, 13 (21), 14454–14469. 10.1021/ACSCATAL.3C03417.

(19) Mazurenko, S.; Prokop, Z.; Damborsky, J. Machine Learning in Enzyme Engineering. ACS Catal. 2020, 10 (2), 1210–1223. 10.1021/ACSCATAL.9B04321.

(20) Kouba, P.; Kohout, P.; Haddadi, F.; Bushuiev, A.; Samusevich, R.; Sedlar, J.; Damborsky, J.; Pluskal, T.; Sivic, J.; Mazurenko, S. Machine Learning-Guided Protein Engineering. ACS Catal. 2023, 13 (21), 13863–13895. 10.1021/ACSCATAL.3C02743.

(21) Meier, J.; Rao, R.; Verkuil, R.; Liu, J.; Sercu, T.; Rives, A. Language Models Enable Zero-Shot Prediction of the Effects of Mutations on Protein Function. Adv. Neural Inf. Process. Syst. 2021, 34, 29287–29303.

(22) Notin, P.; Niekerk, L. Van; Kollasch, A. W.; Ritter, D.; Gal, Y.; Marks, D. S. TranceptEVE: Combining Family-Specific and Family-Agnostic Models of Protein Sequences for Improved Fitness Prediction. bioRxiv 2022, 2022.12.07.519495. 10.1101/2022.12.07.519495.

(23) Alley, E. C.; Khimulya, G.; Biswas, S.; AlQuraishi, M.; Church, G. M. Unified Rational Protein Engineering with Sequence-Based Deep Representation Learning. Nat. Methods 2019, 16 (12), 1315–1322. 10.1038/S41592-019-0598-1.

(24) Cheng, P.; Mao, C.; Tang, J.; Yang, S.; Cheng, Y.; Wang, W.; Gu, Q.; Han, W.; Chen, H.; Li, S.; Chen, Y.; Zhou, J.; Li, W.; Pan, A.; Zhao, S.; Huang, X.; Zhu, S.; Zhang, J.; Shu, W.; Wang, S. Zero-Shot Prediction of Mutation Effects with Multimodal Deep Representation Learning Guides Protein Engineering. Cell Research 2024 34:9 2024, 34 (9), 630–647. 10.1038/s41422-024-00989-2.

(25) Shanker, V. R.; Bruun, T. U. J.; Hie, B. L.; Kim, P. S. Unsupervised Evolution of Protein and Antibody Complexes with a Structure-Informed Language Model. Science 2024, 385 (6704), 46–53. 10.1126/SCIENCE.ADK8946.

(26) Cheng, J.; Novati, G.; Pan, J.; Bycroft, C.; Žemgulyte, A.; Applebaum, T.; Pritzel, A.; Wong, L. H.; Zielinski, M.; Sargeant, T.; Schneider, R. G.; Senior, A. W.; Jumper, J.; Hassabis, D.; Kohli, P.; Avsec, Ž. Accurate Proteome-Wide Missense Variant Effect Prediction with AlphaMissense. Science (1979). 2023, 381 (6664). 10.1126/SCIENCE.ADG7492.

(27) Brandes, N.; Goldman, G.; Wang, C. H.; Ye, C. J.; Ntranos, V. Genome-Wide Prediction of Disease Variant Effects with a Deep Protein Language Model. Nat. Genet. 2023, 55 (9), 1512–1522. 10.1038/S41588-023-01465-0.

(28) Lin, Z.; Akin, H.; Rao, R.; Hie, B.; Zhu, Z.; Lu, W.; Smetanin, N.; Verkuil, R.; Kabeli, O.; Shmueli, Y.; dos Santos Costa, A.; Fazel-Zarandi, M.; Sercu, T.; Candido, S.; Rives, A. Evolutionary-Scale Prediction of Atomic-Level Protein Structure with a Language Model. Science (1979). 2023, 379 (6637), 1123–1130. 10.1126/SCIENCE.ADE2574.

(29) Harding-Larsen, D.; Lax, B. M.; García, M. E.; Mendonça, C.; Mejia-Otalvaro, F.; Welner, D. H.; Mazurenko, S. PRIZM: Combining Low-N Data and Zero-Shot Models to Design Enhanced Protein Variants. bioRxiv 2026, 2026.02.08.704628. 10.64898/2026.02.08.704628.

(30) Hsu, C.; Verkuil, R.; Liu, J.; Lin, Z.; Hie, B.; Sercu, T.; Lerer, A.; Rives, A. Learning Inverse Folding from Millions of Predicted Structures. bioRxiv 2022, 2022.04.10.487779. 10.1101/2022.04.10.487779.

(31) Rives, A.; Meier, J.; Sercu, T.; Goyal, S.; Lin, Z.; Liu, J.; Guo, D.; Ott, M.; Zitnick, C. L.; Ma, J.; Fergus, R. Biological Structure and Function Emerge from Scaling Unsupervised Learning to 250 Million Protein Sequences. bioRxiv 2020, 622803. 10.1101/622803.

(32) Zhu, X.; Robinson, D. A.; McEwan, A. R.; O’Hagan, D.; Naismith, J. H. Mechanism of Enzymatic Fluorination in Streptomy es Cattleya. J. Am. Chem. Soc. 2007, 129 (47), 14597–14604. 10.1021/JA0731569.

(33) Abramson, J.; Adler, J.; Dunger, J.; Evans, R.; Green, T.; Pritzel, A.; Ronneberger, O.; Willmore, L.; Ballard, A. J.; Bambrick, J.; Bodenstein, S. W.; Evans, D. A.; Hung, C. C.; O’Neill, M.; Reiman, D.; Tunyasuvunakool, K.; Wu, Z.; Žemgulytė, A.; Arvaniti, E.; Beattie, C.; Bertolli, O.; Bridgland, A.; Cherepanov, A.; Congreve, M.; Cowen-Rivers, A. I.; Cowie, A.; Figurnov, M.; Fuchs, F. B.; Gladman, H.; Jain, R.; Khan, Y. A.; Low, C. M. R.; Perlin, K.; Potapenko, A.; Savy, P.; Singh, S.; Stecula, A.; Thillaisundaram, A.; Tong, C.; Yakneen, S.; Zhong, E. D.; Zielinski, M.; Žídek, A.; Bapst, V.; Kohli, P.; Jaderberg, M.; Hassabis, D.; Jumper, J. M. Accurate Structure Prediction of Biomolecular Interactions with AlphaFold 3. Nature 2024, 630 (8016), 493–500. 10.1038/S41586-024-07487-W.

(34) Hopf, T. A.; Green, A. G.; Schubert, B.; Mersmann, S.; Schärfe, C. P. I.; Ingraham, J. B.; Toth-Petroczy, A.; Brock, K.; Riesselman, A. J.; Palmedo, P.; Kang, C.; Sheridan, R.; Draizen, E. J.; Dallago, C.; Sander, C.; Marks, D. S. The EVcouplings Python Framework for Coevolutionary Sequence Analysis. Bioinformatics 2019, 35 (9), 1582–1584. 10.1093/BIOINFORMATICS/BTY862.

(35) Genee, H. J.; Bonde, M. T.; Bagger, F. O.; Jespersen, J. B.; Sommer, M. O. A.; Wernersson, R.; Olsen, L. R. Software-Supported USER Cloning Strategies for Site-Directed Mutagenesis and DNA Assembly. ACS Synth. Biol. 2014, 4 (3), 342–349. 10.1021/SB500194Z.

(36) NEBaseChanger. https://nebasechanger.neb.com/ (accessed 2025-06-23).

