## Supplementary Figures and Tables for "Data-efficient distal engineering of fluorinase using zero-shot models"

SUPPORTING FIGURES

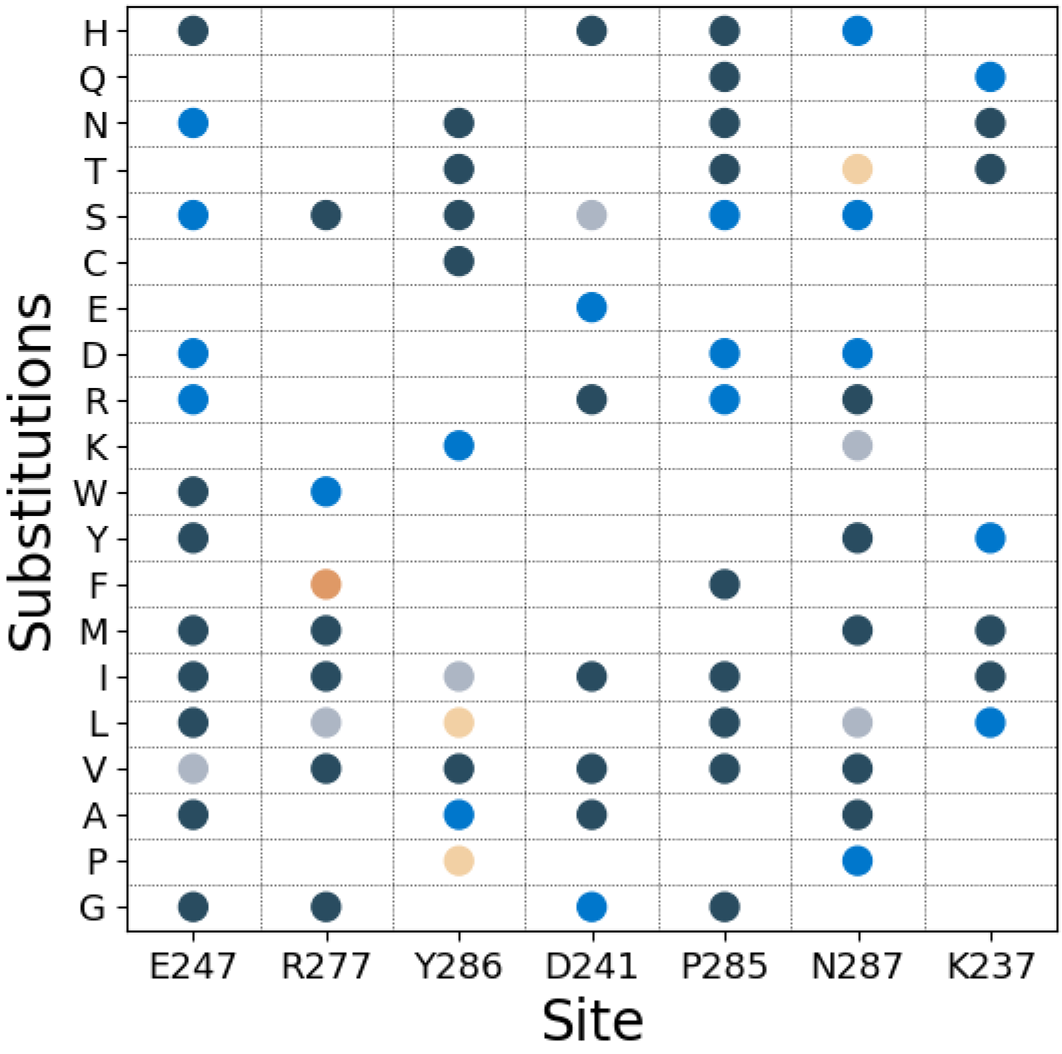


**Figure S1.** Distribution of mutations across the seven hexameric interface residues contained in the FlA low-N dataset (N = 40). Number of occurrences are denoted by color: 1 = dark blue, 2 = blue, 3 = grey, 4 = beige, 5 = orange.

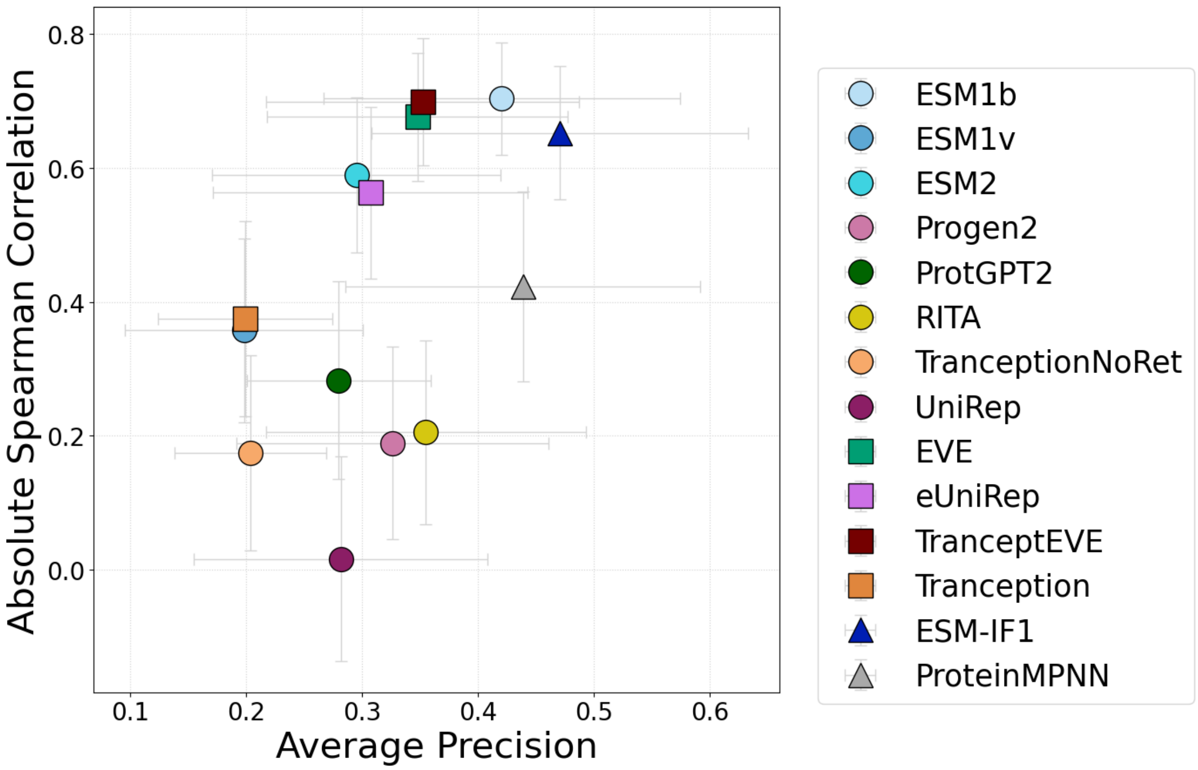


**Figure S2**. Spearman correlation and average precision for zero-shot scores and experimental values for the different zero-shot models of PRIZM (circle = sequence-based model, square = MSA-based model, and triangle = structure-based model). Error bars indicate the standard deviation estimated from bootstrapping the dataset (N = 1000).

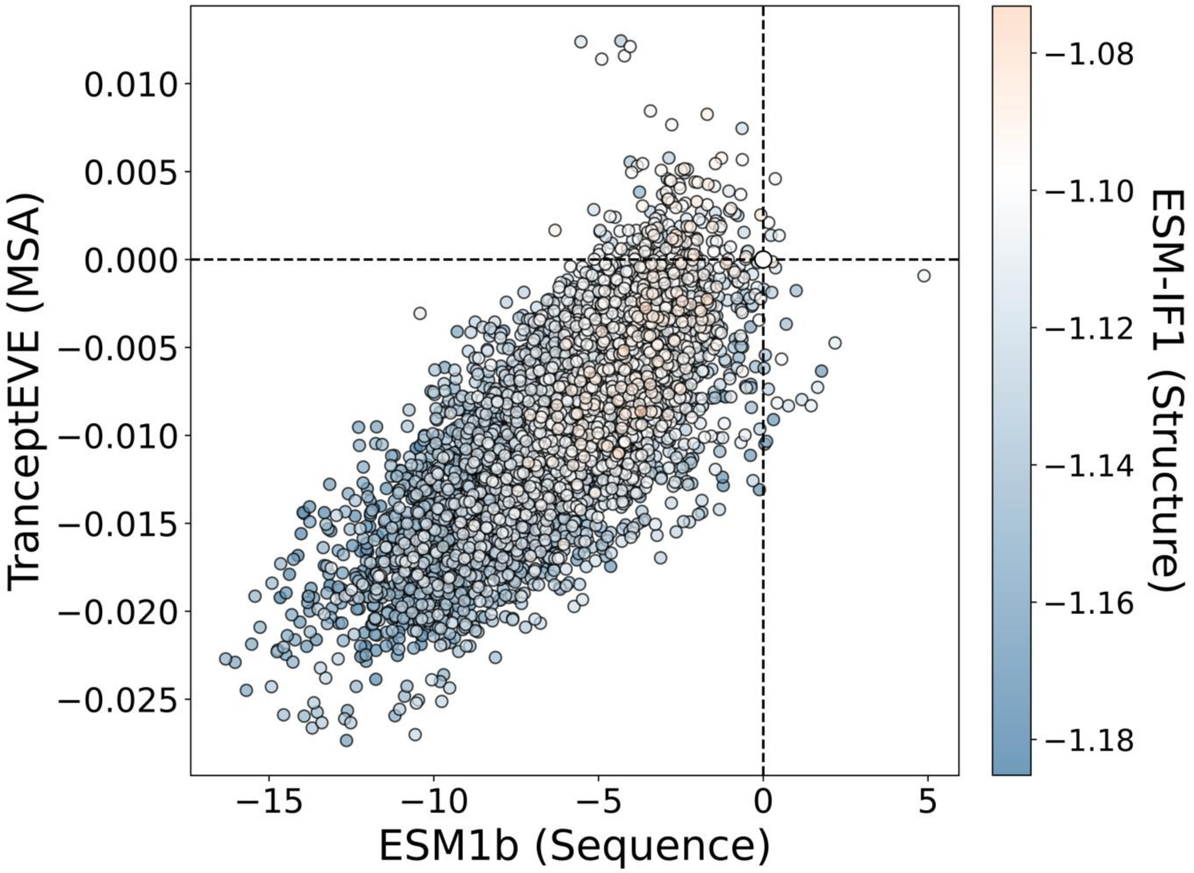


**Figure S3**. Zero-shot scores of all single-point variants using the best models for each category identified by PRIZM using the FlA low-N dataset. WT is denoted by dashed line and white color, while variants predicted by ESM-IF1 to be better or worse than WT are colored orange or blue, respectively.

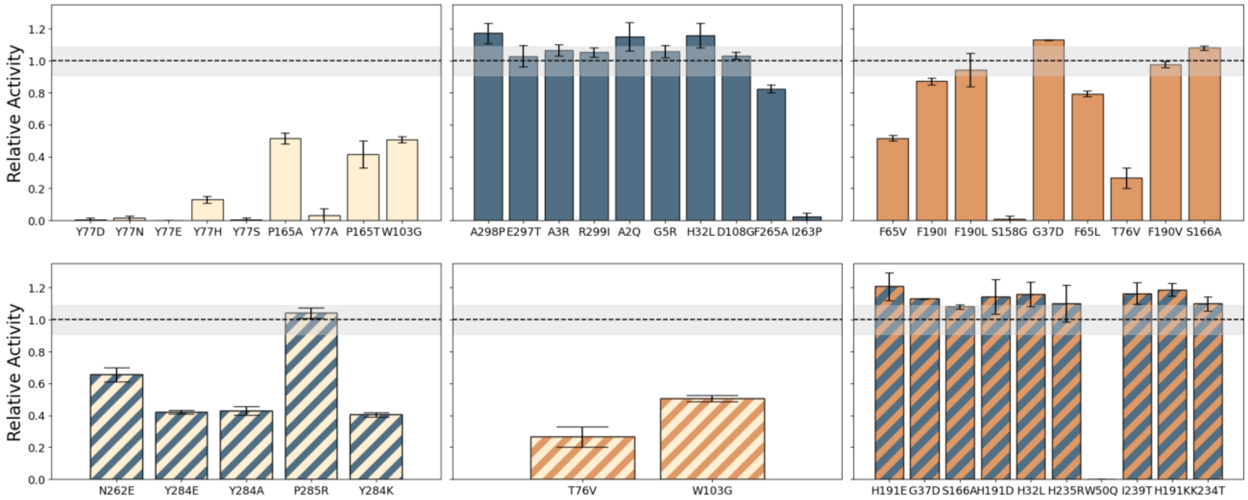


**Figure S4**. Relative activity after two minutes of single mutants selected using the PRIZM workflow. The top row shows variants selected using a greedy top 10 approach for ESM-1b (beige), TranceptEVE (blue), and ESM-IF1 (orange), while the bottom row shows the results of single mutants predicted to be better than WT by two models, with color scheme consistent with individual models. Dashed lines denote the WT value of 1. Error bars indicate standard deviation of biological replicates (N=3).

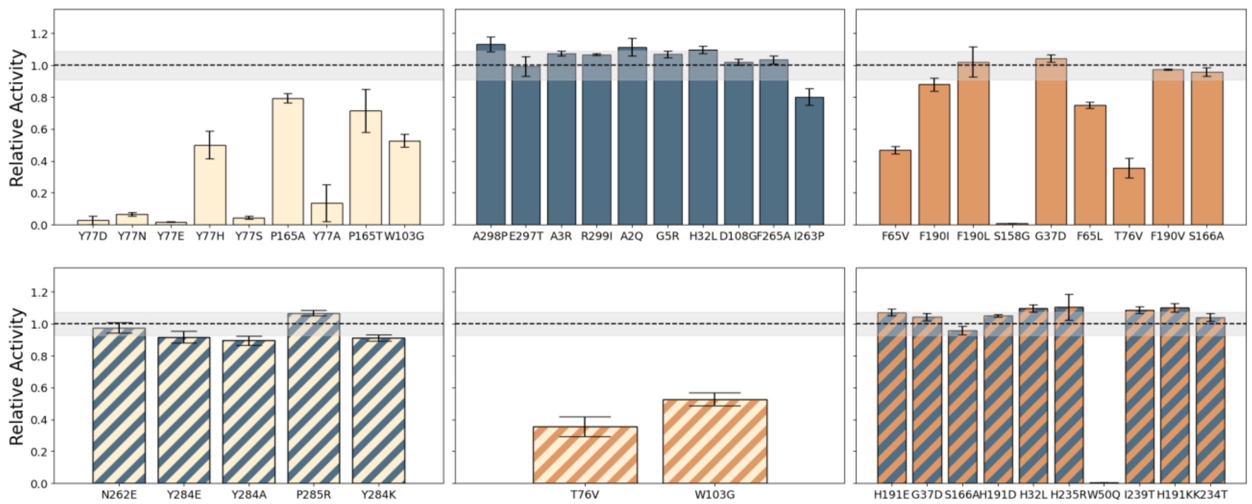


**Figure S5**. Relative activity after three hours of single mutants selected using the PRIZM workflow. The top row shows variants selected using a greedy top 10 approach for ESM-1b (beige), TranceptEVE (blue), and ESM-IF1 (orange), while the bottom row shows the results of single mutants predicted to be better than WT by two models, with color scheme consistent with individual models. Dashed lines denote the WT value of 1. Error bars indicate standard deviation of biological replicates.

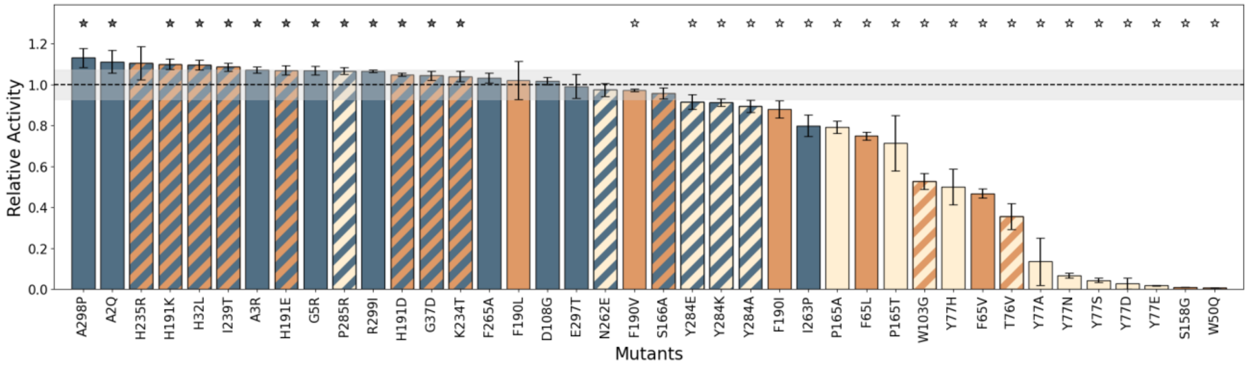


**Figure S6**. Relative activities after three hours of single mutants from different selection criteria: beige = ESM-1b, blue = TranceptEVE, and orange = ESM-IF1. Striped colors denote variants chosen based on multiple models. Although certain variants were selected based on multiple selection criteria, each is displayed only once in this figure (for a comprehensive overview of all selections, refer to Figure S5). The dashed line denote the WT value of 1. Error bars indicate standard deviation of biological replicates. Stars denote variants with relative activity at two minutes significantly different from WT (one-sided Welch test with α = 0.05, grey = better than WT, white = worse than WT).

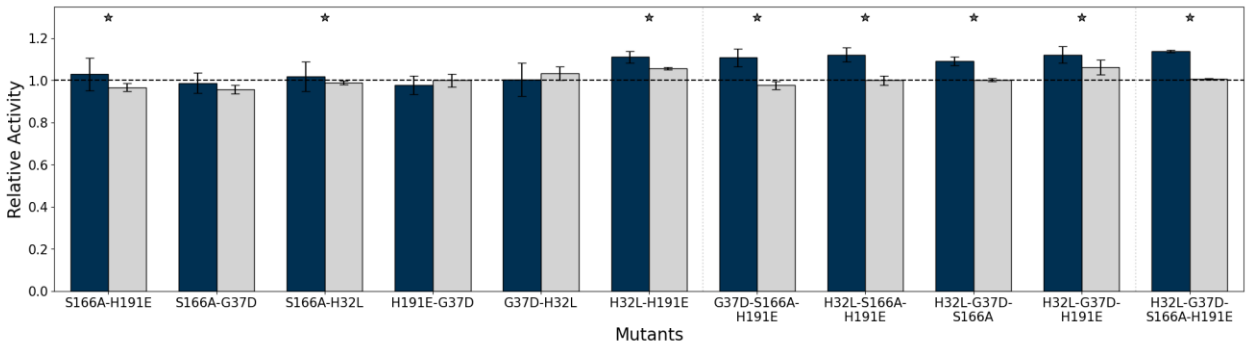


**Figure S7**. Relative activities of combinatorial mutants after two minutes (blue) and three hours (grey). The dashed line denotes the WT value of 1. Error bars indicate standard deviation of biological replicates. Variants with relative activity significantly above WT at two minutes (one-sided t-test with $\alpha = 0.05$) are denoted by a star.

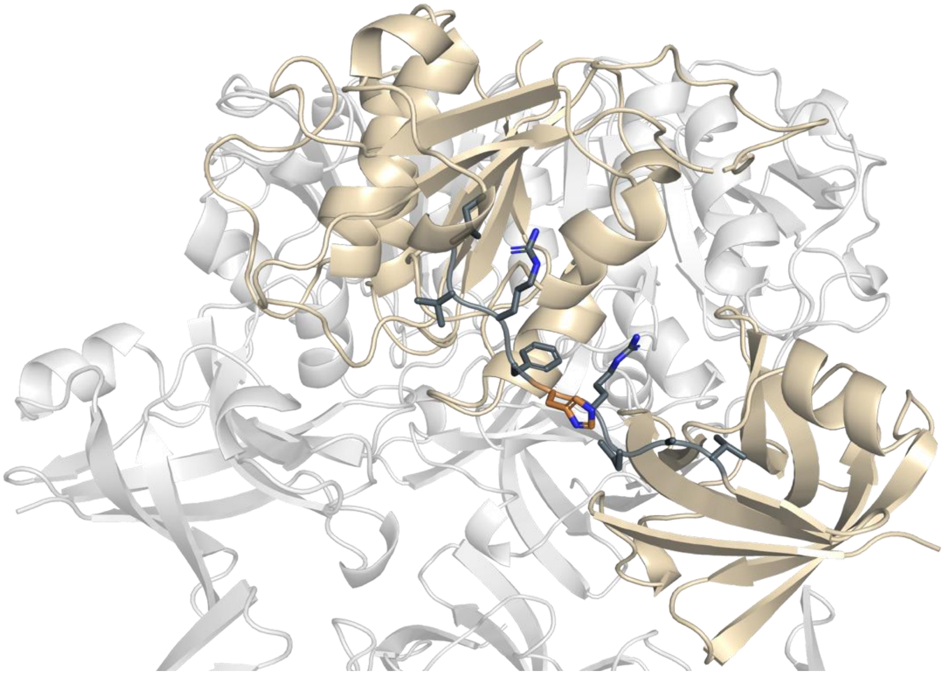

**Figure S8**. The FlA crystal structure (PDB ID: 5B6I) with the monomeric unit colored beige, while the linker region between monomer domains and the H191 residue are shown as sticks and colored grey and orange, respectively.

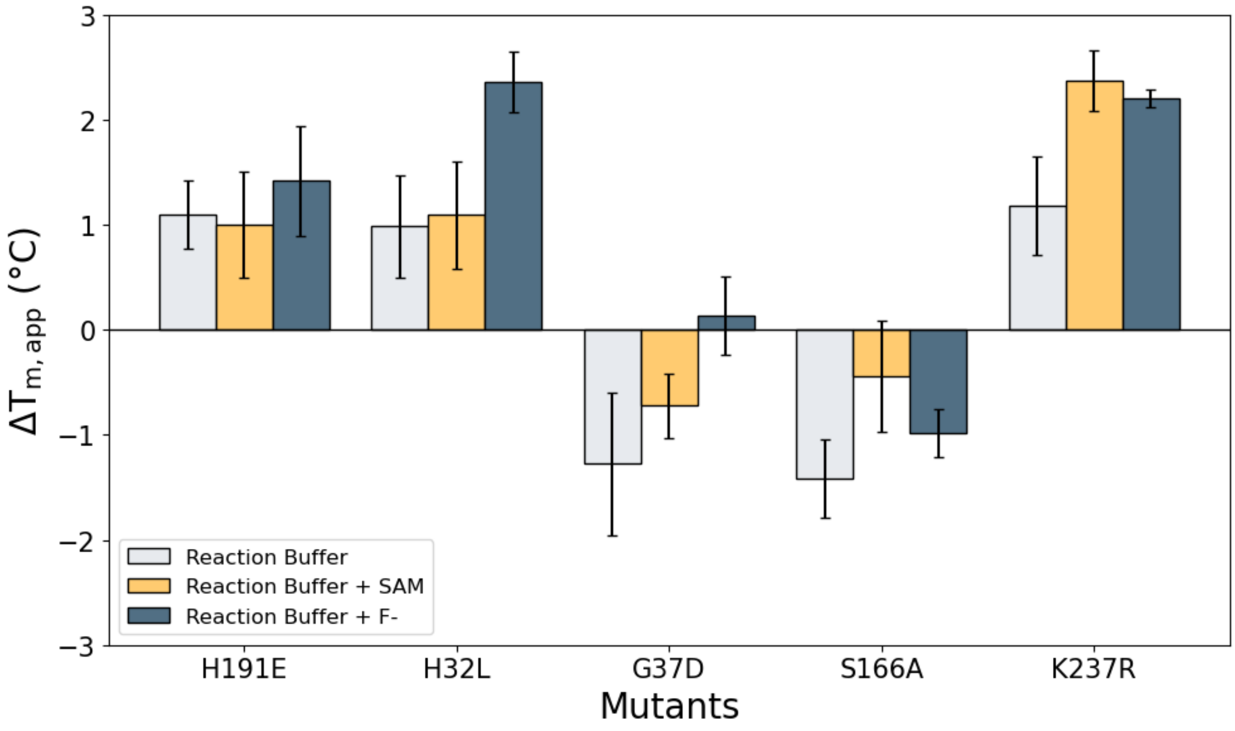


**Figure S9**. $\Delta T_{m,app}$ of WT and top FlA variants at different conditions: reaction buffer alone (grey), with SAM (yellow), or with $F^-$ (blue). Error bars indicate the standard deviation of biological replicates with N = 3.

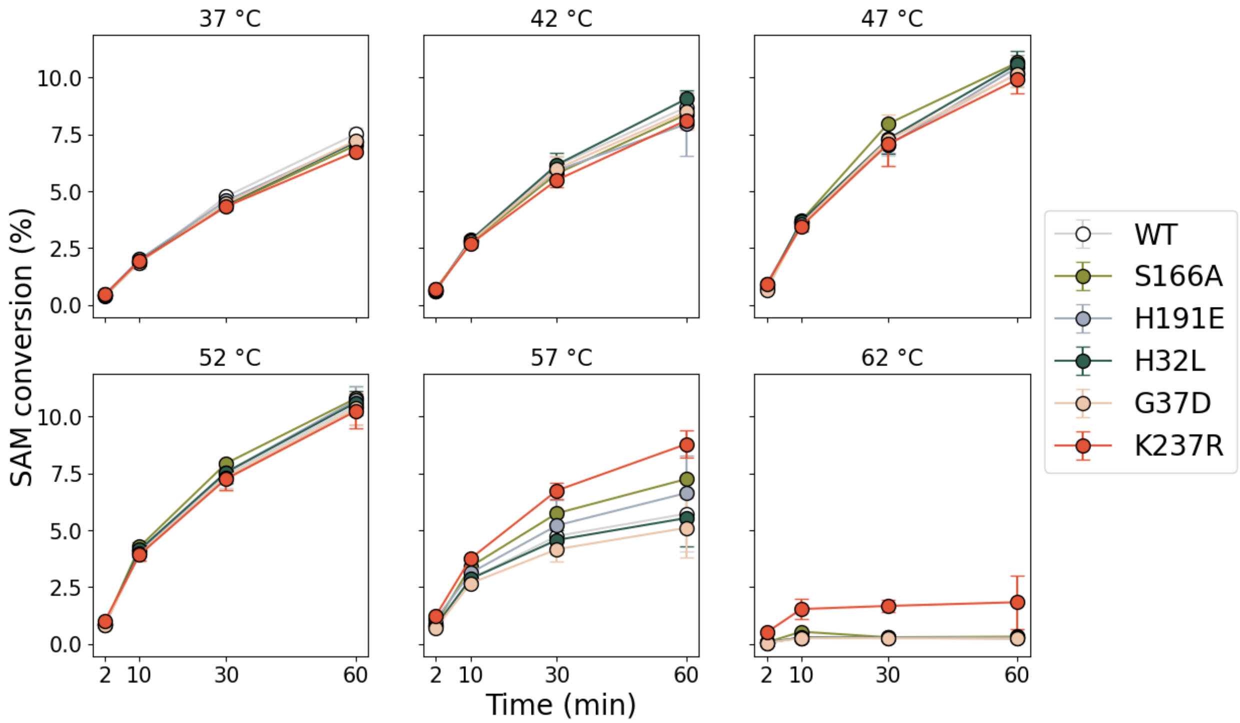


**Figure S10**. SAM conversion (%) for WT and top FlA variants at different temperatures and time points. Error bars indicate the standard deviation of biological replicates with N = 3.

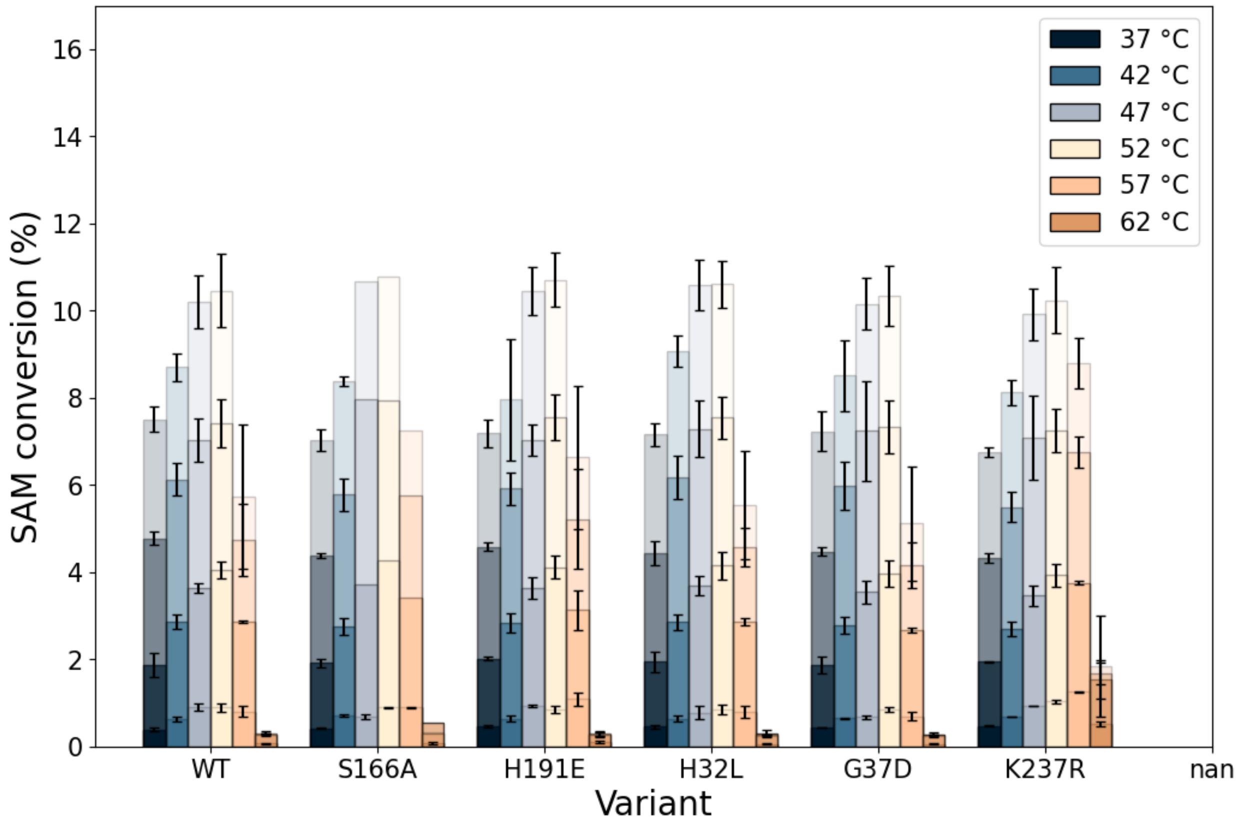


**Figure S11**. SAM conversion (%) for WT and top FlA variants at different temperatures and time points. Error bars indicate the standard deviation of biological replicates with N = 3.

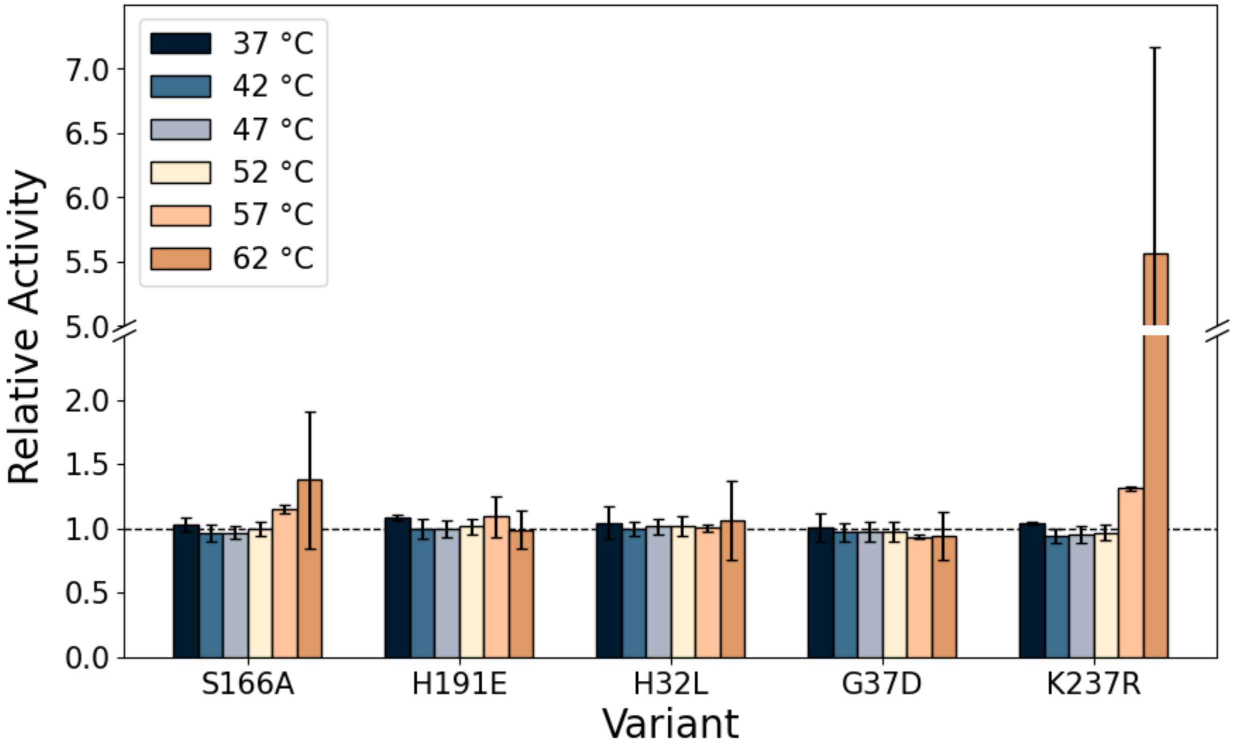


**Figure S12**. Relative activity of WT FlA and top FlA after ten minutes at different temperatures. Error bars indicate the standard deviation of biological replicates with N = 3, while the dashed line indicates the relative activity of 1.

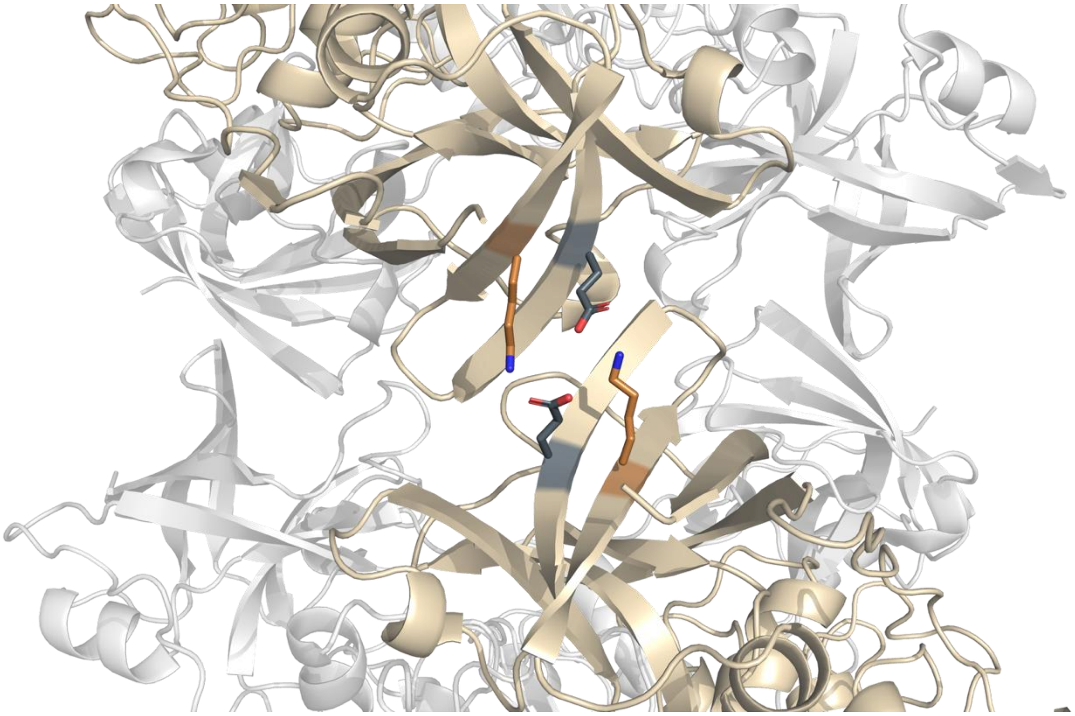

**Figure S13**. The FlA crystal structure (PDB ID: 5B6I) with two interacting C-terminal (hexameric interface) domains colored beige; the K237 and E247 residues are shown as sticks and colored orange and grey, respectively.

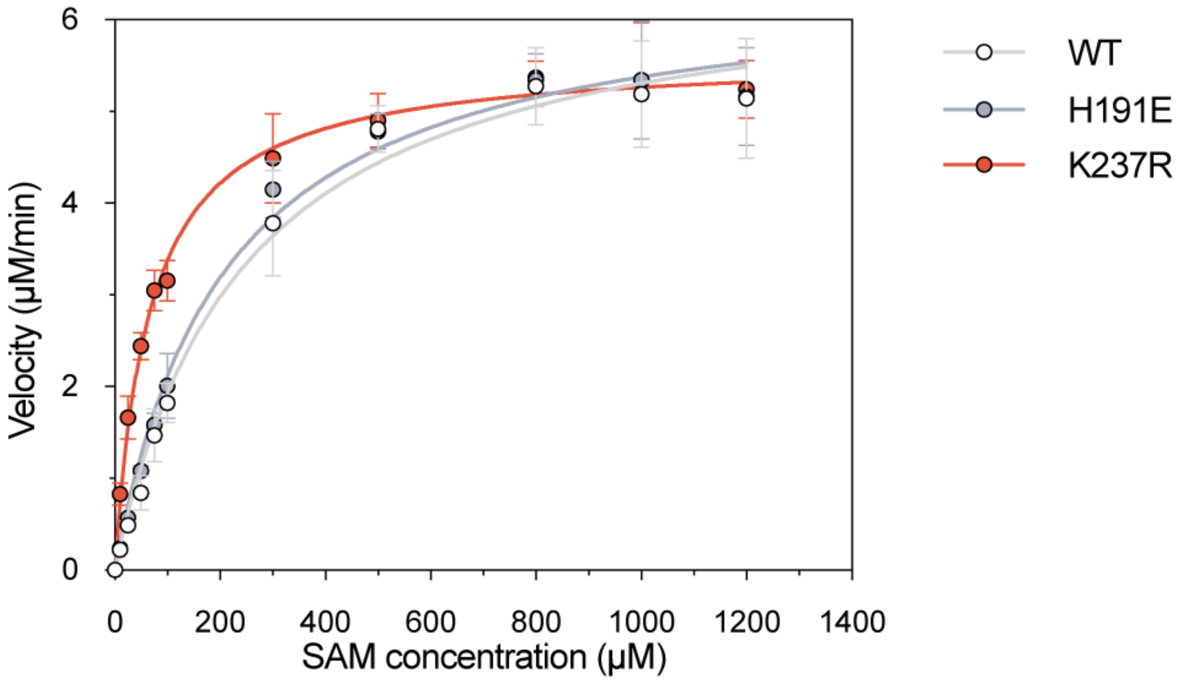


**Figure S14**. Michaelis-Menten kinetics of WT, H191E, and K237R measured across increasing SAM concentrations. Points show mean initial velocities, error bars indicate the standard deviation of biological replicates with N = 3, and curves are nonlinear Michaelis-Menten fits using GraphPad Prism version 10.6.1.

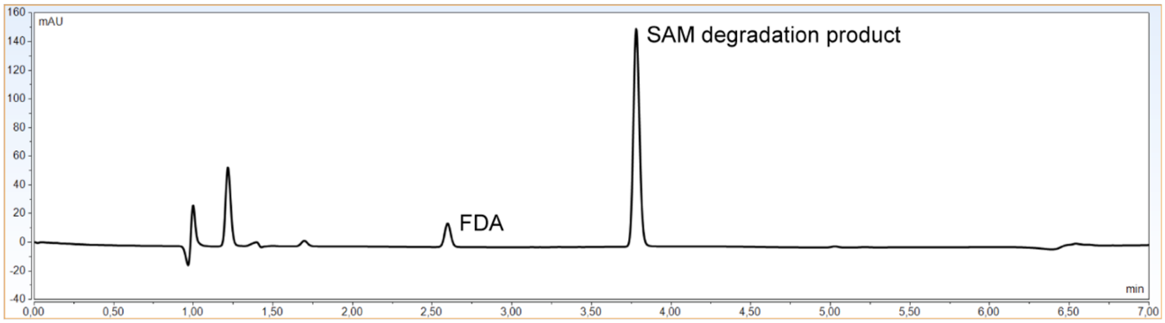


**Figure S15**. Representative HPLC trace showing FDA and SAM degradation product peaks at 260 nm.

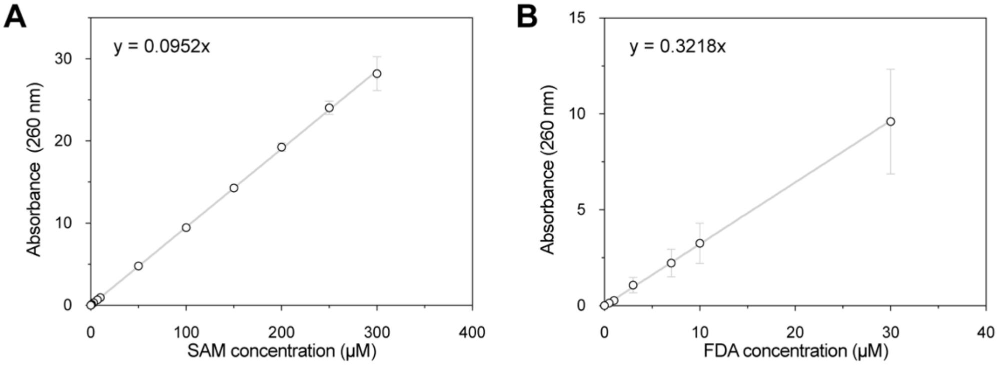


**Figure S16**. Calibration curves at $A_{260}$ for **A** SAM (0–300 µM) and **B** FDA (0–30 µM). Points show mean absorbance, error bars indicate the standard deviation of biological replicates with N = 3, and lines indicate linear fits (slopes denoted on plots).

SUPPPORTING TABLES

**Table S1.** FIA variants from the original low-N dataset, with measured relative activity compared to wildtype (set at 100%).

| Mutant ID | Mutations | Relative Activity (%) |
|---|---|---|
| HiM_001 | E247G, R277L, Y286K | 16.31 |
| HiM_002 | D241G, E247V | 31.38 |
| HiM_003 | P285R, N287P | 12.97 |
| HiM_004 | K237T, E247D, Y286P | 4.60 |
| HiM_005 | P285V, Y286A, N287R | 2.40 |
| HiM_007 | P285G, N287L | 1.71 |
| HiM_010 | E247S | 99.43 |
| HiM_012 | D241G, E247M, N287L | 0.00 |
| HiM_013 | E247V, R277F, N287K | 0.00 |
| HiM_014 | E247N, R277G, Y286N | 0.00 |
| HiM_015 | E247D | 9.39 |
| HiM_016 | E247Y, P285D | 96.00 |
| HiM_019 | D241E, E247L, Y286I | 0.00 |
| HiM_020 | Y286P, N287H | 0.00 |
| HiM_021 | R277W, Y286S, N287K | 0.00 |
| HiM_027 | R277F, N287L | 0.00 |
| HiM_029 | P285L, Y286I | 75.29 |
| HiM_030 | K237Q, D241E, R277F, P285Q, Y286I | 0.00 |
| HiM_031 | N287T | 48.85 |
| HiM_032 | K237M, D241I, P285S, N287A | 0.00 |
| HiM_033 | K237I, E247I, R277V, Y286T, N287S | 0.00 |
| HiM_034 | K237N, E247N, Y286L, N287T | 1.72 |
| HiM_035 | N287Y | 95.98 |
| HiM_036 | D241H, R277W, P285N | 0.00 |
| HiM_037 | K237Y, R277M, P285S | 18.39 |
| HiM_038 | K237L, E247R, Y286P, N287S | 0.00 |
| HiM_039 | D241S, N287M | 0.57 |
| HiM_040 | Y286L, N287V | 5.75 |
| HiM_041 | D241V, E247V, R277S, Y286L, N287P | 0.00 |
| HiM_042 | Y286V, N287D | 3.84 |
| HiM_043 | K237Q, D241R, E247W, P285F, Y286A | 0.00 |

| | | |
|---|---|---|
| HiM_044 | D241S, N287H | 80.40 |
| HiM_045 | P285R, N287T | 94.89 |
| HiM_046 | E247H, R277F, N287K | 87.22 |
| HiM_047 | E247R, R277F, Y286P, N287D | 0.00 |
| HiM_048 | P285T, N287T | 0.00 |
| HiM_049 | K237Y, R277L | 1.14 |
| HiM_050 | E247A, R277I, P285I, Y286K | 0.00 |
| HiM_051 | K237L, D241A, P285D, Y286C | 1.70 |
| HiM_052 | D241S, E247S, R277L, P285H, Y286L | 0.00 |

**Table S2**. FlA PRIZM variants, with measured relative activity compared to wildtype (set at 100%) and standard deviation of biological replicates (N = 3).

| Category | Mutations[a] | Relative Activity at 2 min (%)[b] | Relative Activity at 180 min (%)[b] |
|---|---|---|---|
| *ESM-1b Rankings* | | | |
| 1 | Y77D | 0.55 ± 0.95 | 2.82 ± 2.76 |
| 2 | Y77N | 1.48 ± 1.31 | 6.76 ± 1.14 |
| 3 | P165N | N/A | N/A |
| 4 | Y77E | 0.00 ± 0.00 | 0.00 ± 0.00 |
| 5 | Y77H | 13.05 ± 2.06 | 49.93 ± 8.70 |
| 6 | Y77S | 0.63 ± 1.10 | 4.45 ± 0.95 |
| 7 | P165A | 51.94 ± 3.47 | 79.23 ± 2.92 |
| 8 | Y77A | 3.29 ± 4.24 | 13.54 ± 11.67 |
| 9 | P165T | 41.47 ± 8.52 | 71.50 ± 13.50 |
| 10 | W103G | 50.70 ± 2.02 | 52.67 ± 3.95 |
| *TranceptEVE Rankings* | | | |
| 1 | A298P | 117.22 ± 6.39 | 113.16 ± 4.69 |
| 2 | E297T | 102.89 ± 6.58 | 99.23 ± 6.00 |
| 3 | A3R | 106.65 ± 3.46 | 107.26 ± 1.44 |
| 4 | R299I | 105.24 ± 2.93 | 106.59 ± 0.58 |
| 5 | A2Q | 115.00 ± 8.76 | 111.27 ± 5.52 |
| 6 | G5R | 105.80 ± 4.04 | 106.59 ± 2.09 |
| 7 | **H32L** | 115.68 ± 7.67 | 109.61 ± 2.34 |
| 8 | D108G | 103.20 ± 2.04 | 101.95 ± 1.81 |
| 9 | F265A | 82.38 ± 2.55 | 103.37 ± 2.44 |
| 10 | I263P | 2.52 ± 2.27 | 79.97 ± 5.21 |
| *ESM-IF1 Rankings* | | | |
| 1 | F65V | 51.51 ± 1.81 | 46.79 ± 2.22 |
| 2 | F65I | N/A | N/A |
| 3 | F190I | 87.09 ± 2.16 | 87.89 ± 4.19 |
| 4 | F190L | 94.11 ± 10.41 | 102.07 ± 9.39 |
| 5 | S158G | 1.03 ± 1.79 | 0.92 ± 0.055 |
| 6 | **G37D** | 112.97 ± 0.07 | 104.40 ± 2.35 |
| 7 | F65L | 79.30 ± 1.77 | 74.91 ± 1.90 |

| | | | |
|---|---|---|---|
| 8 | T76V | 26.69 ± 6.39 | 35.61 ± 6.26 |
| 9 | F190V | 97.75 ± 2.05 | 97.33 ± 0.54 |
| 10 | **S166A** | 108.04 ± 1.34 | 95.79 ± 2.78 |
| *ESM-1b & TranceptEVE* | | | |
| Above WT | N262E | 65.52 ± 4.31 | 97.45 ± 3.21 |
| Above WT | Y284E | 42.14 ± 0.99 | 91.69 ± 3.59 |
| Above WT | P285R | 104.09 ± 3.27 | 106.73 ± 1.76 |
| Above WT | Y284A | 43.02 ± 2.77 | 89.48 ± 2.95 |
| Above WT | Y284K | 40.45 ± 1.24 | 91.17 ± 1.84 |
| *ESM-1b & ESM-IF1* | | | |
| Above WT | T76V | 26.69 ± 6.39 | 35.61 ± 6.26 |
| Above WT | W103G | 50.70 ± 2.02 | 52.67 ± 3.95 |
| *TranceptEVE & ESM-IF1*[c] | | | |
| 1 | **H191E** | 120.73 ± 8.59 | 107.05 ± 2.17 |
| 2 | **G37D** | 112.97 ± 0.07 | 104.40 ± 2.35 |
| 3 | **S166A** | 108.04 ± 1.34 | 95.79 ± 2.78 |
| 4 | H191D | 114.36 ± 10.90 | 104.92 ± 0.77 |
| 5 | **H32L** | 115.68 ± 7.67 | 109.61 ± 2.34 |
| 6 | H235R | 109.93 ± 11.60 | 110.53 ± 8.18 |
| 7 | W50Q | 0.00 ± 0.00 | 0.62 ± 0.02 |
| 8 | I239T | 116.34 ± 6.68 | 108.59 ± 2.04 |
| 9 | H191K | 118.70 ± 4.06 | 110.01 ± 2.59 |
| 10 | K234T | 109.86 ± 4.47 | 103.93 ± 2.53 |

[a] Bold names denote variants chosen for combinatorial analysis. [b] N/A: variants unable to express.

[c] Ranking based on combined ranking of the two models.

**Table S3**. MSA of fluorinase homologues[1] colored by PRIZM predictions[a]. Only positions with PRIZM predictions shown.

| ResID[b] | MA37 | PtaU1[c] | SAJ15[c] | Sxin[c] | Stro | Tnor | Pbac | Cbac | Scat | Amza | N902 | Nbra2 | Nbra3 | Nbra1 | AN11 | Abar | CA12 | PRIZM[d] |
|---|---|---|---|---|---|---|---|---|---|---|---|---|---|---|---|---|---|---|
| 2 | A | R | T | S | - | - | N | T | A | M | - | T | T | T | A | - | - | S |
| 3 | A | D | S | A | - | - | K | S | A | S | M | T | T | T | S | M | M | R |
| 5 | G | G | G | P | - | M | V | P | S | S | A | N | N | N | R | T | K | QNTRK |
| 17 | L | L | L | L | V | L | L | L | L | L | L | L | L | L | L | L | L | F |
| 26 | Q | I | Q | Q | L | L | I | I | Q | Q | Q | Q | Q | Q | Q | Q | Q | I |
| 32 | H | L | L | H | Y | I | L | L | Y | M | L | L | L | L | L | L | L | L |
| 37 | G | D | D | G | A | E | E | D | D | D | G | D | D | D | D | D | D | ED |
| 50 | W | W | F | W | F | F | F | F | W | W | W | W | W | W | W | W | W | FQ |
| 83 | T | S | E | E | A | E | M | S | T | T | A | T | T | T | T | G | T | AS |
| 92 | R | - | R | K | - | K | K | R | K | K | K | A | A | A | K | K | K | K |
| 106 | S | - | S | S | - | P | A | A | S | S | S | S | S | S | S | S | S | A |
| 108 | D | - | A | G | - | G | F | E | A | E | A | A | A | A | A | A | A | PAE |
| 110 | F | - | F | F | - | M | I | I | F | F | F | F | F | F | F | F | F | IE |
| 114 | D | P | E | E | E | E | V | E | E | E | E | E | E | E | E | E | E | E |
| 120 | I | V | I | V | V | V | V | I | I | I | I | I | I | I | I | I | I | V |
| 129 | T | R | P | T | F | F | D | S | T | T | T | T | T | T | S | S | S | SLR |
| 137 | I | V | I | I | V | I | L | L | L | T | I | L | L | L | L | L | V | L |
| 146 | K | G | E | K | D | E | N | S | K | D | K | E | E | E | E | E | E | E |
| 166 | S | A | S | A | A | A | S | A | S | S | S | S | S | S | S | S | S | A |
| 176 | A | E | D | S | A | S | S | S | S | E | N | E | E | E | N | E | E | E |
| 181 | R | P | P | P | R | P | P | P | P | K | A | R | R | R | P | K | P | K |
| 183 | D | N | K | E | D | T | G | Q | E | Q | S | A | A | A | A | A | A | A |
| 191 | H | D | S | Q | P | K | K | D | N | T | A | E | E | E | T | D | E | RNPTSAKDE |
| 200 | E | G | T | D | G | N | S | G | E | G | G | N | T | H | G | G | G | SND |
| 203 | S | Q | T | T | R | R | V | A | V | S | S | L | V | V | V | V | V | T |
| 216 | I | I | I | V | V | V | V | V | V | I | L | V | V | V | I | V | V | V |
| 232 | Q | Y | Y | Y | E | Y | Y | Y | Y | Y | Y | Y | Y | Y | Y | Y | Y | AEY |
| 234 | K | T | R | K | L | S | S | T | A | T | T | T | T | T | T | T | T | T |
| 235 | H | N | Q | R | E | R | Q | R | R | N | Q | E | K | K | K | E | R | NLVSQETKR |
| 237 | K | R | R | K | K | T | K | R | R | K | R | R | R | R | K | T | R | R |
| 239 | I | V | T | I | E | V | V | T | T | V | L | T | T | T | V | T | T | VT |
| 252 | P | R | P | P | K | R | Q | P | P | P | P | P | P | P | P | P | P | R |
| 262 | N | K | N | G | Q | D | A | A | N | D | D | A | A | A | T | S | T | SA |
| 292 | L | M | L | L | D | M | M | M | M | M | L | I | I | I | I | I | I | M |

[a] Magenta = residues native to FlA (denoted as MA37), light green = residues predicted by PRIZM to be better than WT, dark green = residues both predicted *and* experimentally shown to be faster than WT. [b] Residue indices are based on MA37. [c] Faster than MA37. [d] PRIZM mutants either predicted to be faster (black), shown to be faster (green), or shown to be slower (red) for the given position.

REFERENCES

(1) Pardo, I.; Bednar, D.; Calero, P.; Volke, D. C.; Damborský, J.; Nikel, P. I. A Nonconventional Archaeal Fluorinase Identified by in Silico Mining for Enhanced Fluorine Biocatalysis. *ACS Catal.* **2022**, *12* (11), 6570–6577. https://doi.org/10.1021/ACSCATAL.2C01184.
